# Molecular complex detection in protein interaction networks through reinforcement learning

**DOI:** 10.1101/2022.06.20.496772

**Authors:** Meghana V. Palukuri, Ridhi S. Patil, Edward M. Marcotte

## Abstract

Many, if not most, proteins assemble into higher-order complexes to perform their biological functions. Such protein-protein interactions (PPI) are often experimentally measured for pairs of proteins and summarized in a weighted PPI network, to which community detection algorithms can be applied to define the various higher-order protein complexes. Current methods, which include both unsupervised and supervised approaches, often assume that protein complexes manifest only as dense subgraphs, and in the case of supervised approaches, focus only on learning *which* subgraphs correspond to complexes, not *how* to find them in a network, a task that is currently solved using heuristics. However, learning to walk trajectories on a network with the goal of finding protein complexes lends itself naturally to a reinforcement learning (RL) approach, a strategy that has not been extensively explored for community detection. Here, we evaluated the use of a reinforcement learning pipeline for community detection in weighted protein-protein interaction networks to detect new protein complexes. Using known complexes, the algorithm is trained to calculate the value of different possible subgraph densities in the process of walking on the network to find a protein complex. Then, a distributed prediction algorithm scales the RL pipeline to search for protein complexes on large PPI networks. The reinforcement learning pipeline applied to a human PPI network consisting of 8k proteins and 60k PPI results in 1,157 protein complexes and shows competitive accuracy with improved speed when compared to previous algorithms. We highlight protein complexes harboring minimally characterized proteins including C4orf19, C18orf21, and KIAA1522, suggest TMC04 to be a putative additional subunit of the KICSTOR complex, and confirm the participation of C15orf41 in a higher-order complex with CDAN1, ASF1A, and HIRA by 3D structural modeling. Reinforcement learning offers several distinct advantages for community detection, including scalability and knowledge of the walk trajectories defining those communities.

## Introduction

Protein-protein interactions (PPIs) are essential to nearly all cellular functions and biological processes. From antibodies binding antigens to block infections to protein filaments comprising cellular cytoskeletons, protein interactions are an important organizational principle across biological scales and organisms. Such multi-protein complexes may additionally bind other molecules, such as DNA, RNA, or metabolites, and play critical roles in cellular processes ranging from DNA replication to transcription to multicellular interactions to tissue organization.

As a consequence, a growing variety of experimental techniques have been developed to determine PPIs at large scale, notably including affinity purification / mass spectroscopy (AP/MS), co-fractionation / mass spectrometry (CF/MS), cross-linking / mass spectrometry (XL/MS), proximity labeling, and yeast two-hybrid assays (Y2H), which are collectively reviewed in [1]–[5]. The resulting PPIs define (often weighted) networks of interactions, in which each node represents a protein, an edge represents the interaction confidence, and certain proximal groups of nodes and edges correspond to multiprotein complexes. Importantly, the experimental methods are not completely accurate and suffer both false positive and negative observations. Hence, integrating PPIs across multiple experiments, as for *e.g*. the networks hu.MAP 1.0 [6] and hu.MAP 2.0 [7] that integrate over 9,000 and 15,000 mass spectrometry experiments respectively from AP/MS [8]–[11] and CF/MS data [12]–[15], can help to mitigate the effects of experimental errors. Combining such approaches with algorithms to cluster proteins and identify complexes from the PPI network should result in more accurate determination of protein complexes. Community detection algorithms can be applied to a PPI network to identify its communities, *i.e*., protein complexes [16].

Community detection methods can be unsupervised, *i.e*., not use any information from known communities in a network and instead rely only on the network topology to cluster it into its communities. Currently, existing unsupervised community detection algorithms tend to rely on many assumptions regarding the topological structures of communities. MCODE (Molecular COmplex DEtection) is an unsupervised method of detecting protein complexes running on the assumption that dense regions of a network represent complexes [17]. Another unsupervised algorithm, CMC (cluster-based on maximal cliques), assumes that communities are mainly in the shape of cliques (again, highly dense subgraphs) [18]. This pattern of similar assumptions carries on to other unsupervised methods such as COACH (core-attachment-based method) [19], ClusterONE (clustering with overlapping neighborhood expansion) [20], and GCE (greedy clique expansion) [21], among others.

On the other hand, supervised community detection methods do consider different topological features of communities apart from density, and learn a community fitness function (*i.e*., the probability of being a community) from known complexes using different learning algorithms. One such approach uses a support vector machine (SCI-SVM) and a Bayesian network (SCI-BN) [22]. For both models, subgraphs are represented using 33 features, and a local subgraph growth process is employed starting from a seed node, with the subgraph growth regulated by limited growth rounds, score improvement over iterations, and extent of overlap with other candidate communities. ClusterSS, cluster with supervised and structural information, is a supervised algorithm using a neural network, 17 subgraph features, and a structural scoring function [23]. All three methods, SCI-SVM, SCI-BN, and ClusterSS use a greedy heuristic algorithm for selecting the neighbor to add to the subgraph in the growth process, with ClusterSS considering only the top neighbors by degree for speed improvements. However, since the methods use serial candidate community sampling, this negatively impacts their scalability to large networks like hu.MAP 1.0 [6] with ∼8k proteins and ∼60k interactions, and hu.MAP 2.0 [7] with ∼10k proteins and over 40k interactions. To combat this, Super.Complex (supervised complex detection algorithm) was developed for high scalability and accuracy [24]. By using AutoML (Automated Machine Learning), it explores different supervised algorithms to utilize the optimal one learned from known communities. Then, it samples candidate subgraphs using the learned fitness function by growing them with an epsilon-greedy heuristic, along with one of four additional heuristics (pseudo metropolis, clique - pseudo metropolis, iterative simulated annealing, and greedy). However, the AutoML pipeline can take a long computation time and the method can still be improved in terms of accuracy.

While current supervised learning methods learn community fitness functions, they do not learn trajectories on the network that can lead to a protein complex, potentially missing out on complexes that cannot be traversed using the heuristics employed, for instance in a greedy setting where the community fitness function is maximized at each step. We can apply reinforcement learning to learn paths traversed on a graph to recall known complexes with high accuracy. MARL (Multi-Agent Reinforcement Learning) uses Q-learning to form clusters in networks based on a multi-agent environment [25]. In this algorithm, each node is viewed as an agent and each agent chooses actions to grow into a cluster. A team reward is given to train the action-value function, based on the modularity of the partition, which again assumes that all communities are dense subgraphs. The team reward can also lead to unstable learning of each agent’s behaviors, apart from taking a long time to compute. Nevertheless, MARL has suggested that there is a lot of potential for the use of reinforcement learning in community detection algorithms.

Rather than using a multi-agent approach with a team reward based on modularity, an unsupervised measure, we use a single agent approach (where a subgraph is viewed as an agent) with a supervised reward from training communities based on whether or not the agent selects the right neighbor to grow the subgraph. The rewards are used to train a value iteration algorithm to learn an optimal value function that is used to predict new communities. During the prediction phase, we implement a parallel method with a single agent on each core (process) to increase the speed of the algorithm by predicting communities in a distributed fashion. Applying the reinforcement-learning (RL) algorithm on a human PPI network after learning from experimentally characterized complexes, we can identify candidate protein complexes, and these candidates can then be experimentally characterized. We can create reliable models of protein complexes that allow us to extract more information about their stability, affinity, and specificity.

In the current work, we formulate community detection as a reinforcement learning task, and implement a value iteration algorithm, learning from known communities. The RL algorithm accurately and efficiently predicts candidate complexes by learning and using a value function from known communities, that maps the density of a subgraph to the probability (score) that traversing the subgraph will yield a protein complex. The algorithm trains on known complexes that have nodes and edges from the network, to accurately optimize scores for the various densities that could occur on the network. Then, the RL pipeline uses these scores to traverse the network by starting with different seed nodes to create candidate complexes in parallel. We apply the reinforcement learning algorithm to a human PPI network consisting of 8k proteins and 60k pairwise protein interactions, resulting in 1,157 candidate protein complexes, including complexes containing uncharacterized proteins that may suggest possible functions for these as yet understudied human proteins.

## Materials and methods

### Reinforcement learning

Reinforcement learning (RL) uses machine learning to enable an agent to make a sequence of decisions based on rewarding desired behaviors and punishing undesired ones. These rewards reinforce the right decisions so that the agent repeats them. Over time, the algorithm finds the best possible decision or action to take in each situation. While training the model, once an error has been corrected, there is a low chance of the same error happening again. This unique process specific to RL makes this an intuitive method for community detection as decision-making for long-term results is needed in the process of traversing a network with the final goal of finding a protein complex.

There are 3 main variables in RL: a state, value function, and reward. A state is a “position” the agent is in, in an environment. A value function is a score given to a state that estimates how beneficial it is for the agent to be in that state to achieve the goal. Lastly, a reward is an incentive that tells the agent if the decision made was correct or not. For example, consider a computer playing against a human in a game of tic tac toe. The AI player is an *agent* created to perform certain *actions* on the tic tac toe grid (the *environment*) based on what *state* the environment is in, in real-time. After each action taken, the agent is given a *reward* based on the four possible outcomes of what the *state* could result in: it winning, the opposing player winning, a draw, or continuing the game. If the agent wins, it will be given a positive reward (*i.e*., 1), and if the opposing player wins or if there is a draw – it will be given a negative reward (*i.e*., -1), and if the game continues there will be no reward (*i.e*., 0 or None). The AI player continues to make moves based on what *state* it currently is in with the goal of maximizing its cumulative reward or *return*. This cycle, also known as an *episode*, is terminated when the game ends. While trying to maximize its rewards, the agent will learn the optimal policy (action to be taken in a particular state) and adapt it based on its learned experience encountering various states.

#### Value iteration

The optimal policy at a state is performing an action that takes the agent to the next best possible state that will maximize the probability of achieving the goal. In other words, the optimal policy, out of the states available, moves the agent to the state having the highest value function. The true or optimal value function, *i.e*., a map of the states to their values can be learned using the value iteration algorithm, a classic reinforcement learning method for problems where a model of the environment dynamics is known, usually with a small number of discrete states. The algorithm is a dynamic programming method that solves the Bellman Optimality Equation (**Equation 1**) iteratively, converging to the optimal value function *V\**.

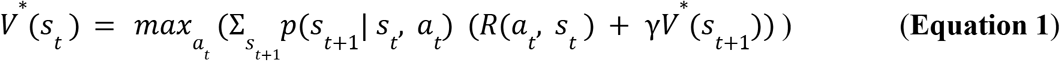

Here, *R* is the defined reward associated with performing action *a*_*t*_ when the state is *s*_*t*_, *γ* is a discount factor and *p* is the transition probability from *s*_*t*_ to *s*_*t+1*_ when it performs action *a*_*t*_.

The value iteration update rule, starting with a value of 0 for all states is given by **Equation 2**:

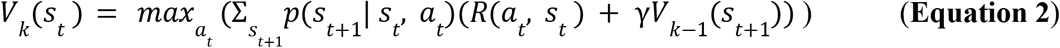

Here, *V*_*k*_ *(s*_*t*_) is the value function of the current state (at time *t* in the episode) in the current iteration *k* (of the value iteration algorithm) and *V*_*k-1*_ *(s*_*t +1*_) is the value function (from the previous iteration *k-1*) of the next possible state.

### Formulating community detection as a reinforcement learning problem

To successfully utilize an RL pipeline for the problem of community detection, the algorithm is first trained on known training communities or complexes. Once the training is deemed to be successful, the learned value function from the training is then used to find complexes on a network.

To learn the value function, each episode consists of starting with a seed node from a training complex and iteratively adding neighbors to grow the subgraph into the complex. This process is then repeated with a new seed node from the training complex. Once all the nodes of the complex have been used as seeds, training moves to the next complex. In this scenario, the agent and environment are defined as the current subgraph in the growth process and the full graph including all of its neighbors, respectively. We represent the state of the agent, *i.e*., the current subgraph by its topological feature, density. The state (density *d*) of the current subgraph is the ratio of the actual number of edges in a subgraph to the total possible number of edges, and can be calculated with **Equation 3**.

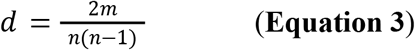

Here *m* is the sum of the edge weights of the edges in the subgraph and *n* is the number of nodes in the subgraph.

For a wide representation of the feature space, the states are discretized into 20 intervals ranging from 0 to 1. The actions performed by the agent comprise adding a neighbor to the current subgraph or terminating the growth process. Choosing a neighboring node in the known complex will provide the agent a positive reward of +0.2, and a negative reward of -0.2 is given if the node chosen is not present in the known complex. The rewards aid the agent in avoiding previous mistakes for it to find an optimal path to create a complex. These rewards allow the state to develop a value function representing the probability of the state resulting in a final community. If none of the remaining neighbors are in the complex, the agent is encouraged to learn to terminate the growth process by receiving a reward of 0, as opposed to choosing a neighbor giving a reward of -0.2.

Once the training completes and an optimal value function is learned, the agent learns candidate complexes on the entire network by starting with seed nodes in parallel and adding neighbors giving the highest value function at each iteration, until the action of terminating a growth process gives a higher value than adding any of the neighbors.

#### Proof of applicability of RL to community detection

The environment is deterministic and the next state the subgraph moves into (*s*_*t+1*_) is only dependent on the previous state (*s*_*t*_), the current subgraph. It does not depend on any other states previously encountered by the agent, satisfying the Markov property (**Equation 4**) with a memoryless process.

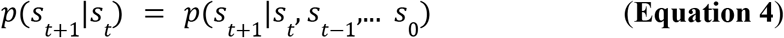

Here, on the left-hand side, *p* is the conditional probability of achieving a state given only the previous state, while on the right-hand side, the probability is conditional on all the previous states encountered. Therefore, with this formulation, community detection can be treated as a Markov Decision Process (MDP) and solved using reinforcement learning methods.

#### Value iteration for community detection

The value iteration update rule with our formulation of the community detection problem is given by **Equation 5**:

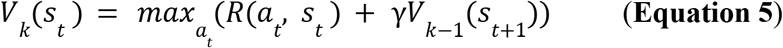

*V*_*k*_ *(s*_*t*_) is the value function of the current state (at time *t* in the episode) in the current iteration *k* (of the value iteration algorithm), *R* is the defined reward, *γ* is the discount factor (0.5), and *V*_*k-1*_ *(s*_*t +1*_) is the value function (from the previous iteration *k-1*) of the next possible state. We obtain this simple update rule, derived from the Bellman optimality equation (**Equation 6**) using a transition probability *p(s*_*t+1*_|*s*_*t*_, *a*_*t*_) of 1 in **Equation 1** due to the deterministic nature of this problem, *i.e*., the state transitions from *s*_*t*_ to only *s*_*t+1*_ when action *a*_*t*_ is taken.

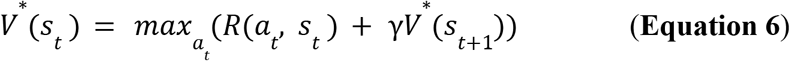

Here, *V*^*^ is the optimal value function, which the algorithm’s value function converges to after a few iterations using the value iteration update rule (**Equation 5**), starting with a value of 0 for all states.

### Reinforcement learning community detection algorithm

#### Overview

There are 3 main steps for community detection on a network using reinforcement learning:

1. **Training** the algorithm to walk across training complexes by learning a value function corresponding to each state (subgraph) encountered in the process.
2. **Finding** candidate complexes by using the learned value function to walk on the human protein-protein interaction network.
3. **Benchmarking** learned complexes against known complexes.

#### Training the algorithm

For training the RL pipeline, we use the known training complexes and represent each complex as a target subgraph of the entire protein-interaction network. The agent, *i.e*., the current subgraph expands by adding, at each step, the neighbor that yields the highest value for the current subgraph (the value is calculated using the reward given for adding this node and the value of the next state, as shown in **Equation 5**). The algorithm updates the value of the density of the current subgraph to this new value (**Equation 5**). Each time a state (density) is encountered in the process of training on multiple training complexes, the value of that state is updated using the update rule (**Equation 5**), moving towards convergence of the value function and eventually learning a value function mapping densities to their probability of leading to a protein complex.

Figure 1. shows an example of learning the value function while traversing a single known complex. The initial subgraph or seed is an edge between an initial random node and the node’s neighbor with whom the node shares an edge with the highest edge weight. To calculate the potential value function of the current subgraph, *i.e*., the term within the max() in **Equation 5** for each neighbor of the current subgraph, each neighbor is temporarily added to the subgraph. The density of that temporary subgraph is calculated, followed by querying the value function for that state along with the reward based on whether or not the neighbor is present in the final protein complex. After calculating the potential value function, the neighbor is removed from the subgraph for this process to be repeated for the rest of the neighbors. Once all of the neighbors have been evaluated, the value function of the current state is updated and the neighbor yielding the state that provided the maximum value function is added to the subgraph. This new subgraph will be the new “seed” as this process repeats itself. The subgraph, or complex, will be “complete” and the algorithm stops adding nodes if all the neighboring nodes return lower value functions than the action of terminating the growth process, represented by adding an imaginary node with reward 0, leading to the same state as before, indicating that no new neighbors should be in the complex. Note how a reward of 0 encourages the algorithm to terminate the growth process when all other options are adding the wrong nodes, *i.e*., choosing actions that have a reward of -0.2. Conversely, if a correct node is available, its reward of 0.2 encourages choosing that node over terminating growth. Once a subgraph is deemed as complete, the process is repeated starting with a different node of the same complex, so that other trajectories to build the same complex are explored and the value function is updated accordingly. After starting with each of the nodes of a complex as seeds, training moves to the next complex, starting with a random new seed from this complex, and the process repeats.

**Figure 1.**
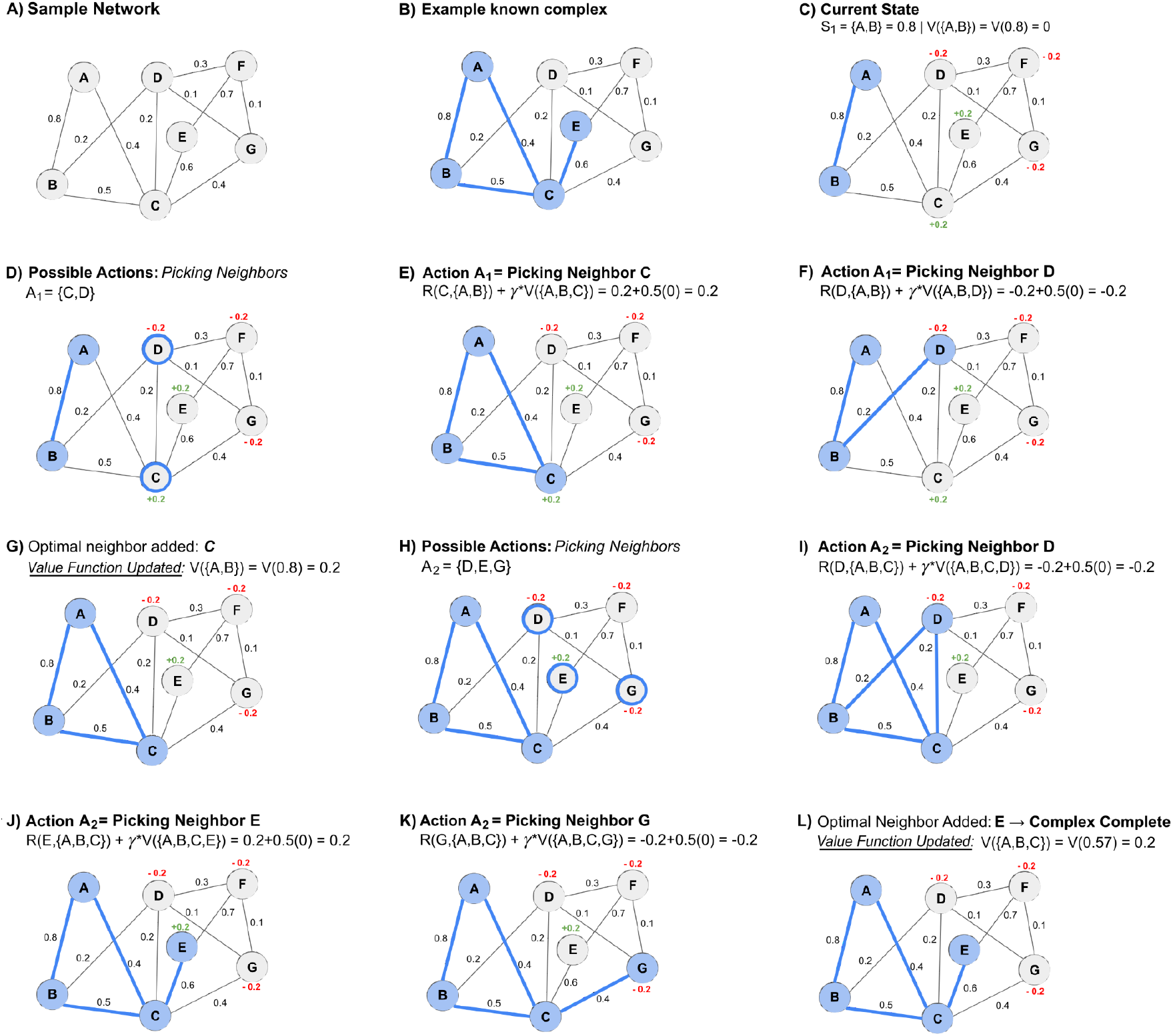
Example trajectory of training the RL pipeline on a sample network by learning a value function. (A)This network comprises 7 nodes and 11 weighted edges **(B)** A known complex consists of the nodes A, B, C, and E. **(C)** First, a seed edge (A, B) is identified to begin the walk. At {A, B}, the state (density) *S*_*1*_ *= 0.8* and the value function *V({A, B}) = V(0.8) = 0* (since values of all densities are initialized to 0). When a node is added to the current subgraph, a reward of +0.2 is given if the node is present in the training complex and a reward of -0.2 is given if absent. **(D)** We evaluate all possible neighbors *i.e*., C and D, to add to the current subgraph {A, B}. Using the value iteration update rule with a discount factor *γ* of 0.5, we add each neighbor as the next state and compute a corresponding value for the current state. **(E)** The temporary complex {A, B, C}, corresponds to *S*_*2*_ *= 0.57* and *V({A, B, C}) = V(0.57) = 0* (since this state has not been encountered before). Using the reward for node C (+0.2), adding node C to the complex {A, B} would give *V({A, B}) = V(0.8) = 0.2*. **(F)** The temporary complex {A, B, D} corresponds to *S*_*2*_ = 0.33, *V({A, B, D}) = V(0.33)* = 0. Using the reward for node D (−0.2), adding node D to the current complex {A, B}, would give *V({A, B}) = V(0.8) = -0.2*. **(G)** Once all neighbors have been evaluated, the neighbor providing the highest value function (C) is added to the candidate complex. The value function of the original state *S*_*1*_ *= {A, B} = 0.8* is updated from 0 to +0.2. **(H)** Again, we evaluate all possible neighbors of the newly updated complex {A, B, C}, *i.e*., D, E, and G, to consider the possible actions the agent can take. **(I)** The temporary complex {A, B, C, D} corresponds to *S*_*3*_ *= 0.35, V({A, B, C, D})* = 0. Using the reward for node D (−0.2), adding node D to the current complex {A, B, C} gives *V({A, B, C}) = V(0.57) = -0.2*. **(J)** The temporary complex {A, B, C, E} corresponds to *S*_*3*_ = 0.38, *V({A, B, C, E})* = 0. Using the reward for node E (+0.2), adding node E to the current complex {A, B, C}, results in *V({A, B, C}) = V(0.57) = 0.2*. **(K)** The temporary complex {A, B, C, G}, corresponds to *S*_*3*_ *= 0.35, V({A, B, C, G})* = 0. Using the reward for node G (−0.2), if we add node G to the current state of the complex {A, B, C}, the resulting value function is -0.2. **(L)** Node E provides the highest value function, so it is the optimal neighbor and is added to the complex. The value function of the complex state *S*_*2*_ *= {A, B, C} = 0.57* is updated from 0 to +0.2. This process of evaluating neighbors is repeated until the action of terminating the growth process is chosen, represented by adding an imaginary node with reward 0, leading to the same state as before. As the remaining neighbors D, F, and G have a reward of -0.2, the imaginary node is chosen as it results in a value function (0.1) higher than the neighbors (D: -0.2, F: -0.2, G: -0.2). At this point, the candidate complex is finalized {A, B, C, E} and the process stops. A new seed edge is chosen from the network and this process repeats, updating the same value function.

#### Finding candidate complexes

Once we observe that the value functions of different states have converged in the training process, we can use the learned value function to walk paths on the network to find complexes.

For each node of the network, we choose its corresponding highest edge-weight as a seed edge to grow a candidate complex. For each seed edge, the neighbors are evaluated and the neighbor yielding the subgraph with the maximum value function is added. The growth process stops when adding any neighbor lowers the value function of the current subgraph. For instance, consider a subgraph with a value function of 0.2. If on evaluating each of the subgraph’s neighbors, each of them returns a value function lesser than 0.2, the algorithm stops and the subgraph will be considered a candidate complex. This process is repeated for the next seed edge in the network. These steps are detailed in **Algorithm 1** and an example of finding a complex on the network is shown in **Figure 2**.

**Figure 2.**
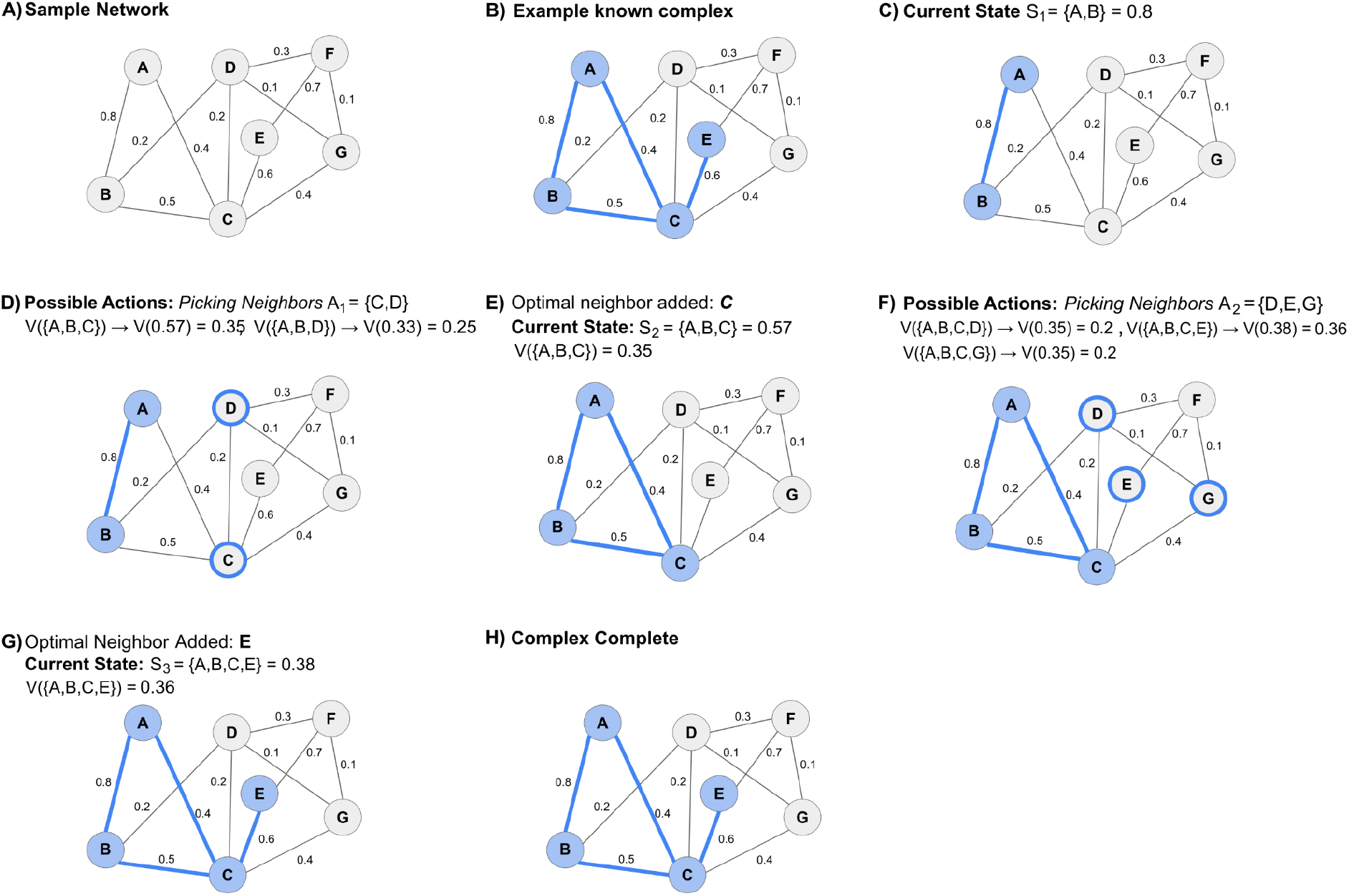
Example trajectory for finding a complex on a sample network with the RL pipeline using a learned value function. **(A)**. This network comprises 7 nodes and 11 edges, with edge weights shown next to each corresponding edge. **(B)** In this network, a known complex consists of the nodes A, B, C, and E. The goal of the algorithm is to predict this known complex from the network using the learned value function. **(C)** The first step is to identify a seed edge to begin the walk (edge AB, with an edge weight of 0.8). At this seed edge, the complex is at state (density) *S*_*1*_ *= 0.8*. **(D)** Then, we evaluate all possible neighbors of nodes A and B, *i.e*., C and D. Adding node C gives a temporary complex {A, B, C} with *S*_*2*_ *= 0.57*, and a learned value *V({A, B, C})* = *V(0.57) = 0.35*, while adding node D gives a temporary complex {A, B, D} with *S*_*2*_ *= 0.33, V({A, B, D})* = *V(0.33) = 0.25*. **(E)** The neighbor with the highest value function is node C and hence, node C is added to the subgraph resulting in *S*_*2*_ *= 0.57, V({A, B, C})* = *V(0.57) = 0.35*. **(F)** The next neighbors of this updated complex are evaluated, *i.e*., D, E, and G. Adding node D leads to *S*_*3*_ *= 0.35, V({A, B, C, D})* = *V(0.35) =* 0.2, node E results in *S*_*3*_ = 0.38, *V({A, B, C, E})* = *V(0.38) = 0.36*, and node G leads to *S*_*3*_ = 0.35, *V({A, B, C, G})* = *V(0.35) = 0.2*. **(G)** Since the neighbor yielding the highest value function is node E, this node is added to the complex resulting in *S*_*3*_ = 0.38, *V({A, B, C, E})* = *V(0.38) = 0.36*. **(H)** Each of the remaining neighbors (D, F, and G) results in a value function less than that of the current complex {A, B, C, E}. Thus, no neighbor is added and the predicted community is complete.

##### Algorithm 1. Finding candidate complexes

For each seed edge in the network:

1. Using the edges of the network, all the neighbors of the seed edge are found.
2. For each neighbor:
  ▪ It is temporarily added to the subgraph.
  ▪ The density (state) is calculated.
  ▪ The corresponding value of the state is obtained from the learned value function and is noted.
  ▪ The neighbor is removed.
3. The neighbor that resulted in the complex having the highest value function is added if this value is greater than the value function of the current subgraph, otherwise, the growth process terminates and the current subgraph is returned as a candidate complex.

#### Post-processing and evaluation

Once we have found all the candidate subgraphs corresponding to the specified seed nodes (in our experiments we use all the nodes of the graph as seed nodes), we perform a post-processing step to merge highly overlapping complexes. Adapting the pairwise merging algorithm in Super.Complex [24], if the overlap of two complexes is more than a specified threshold, we retain the complex with the highest value function of the two complexes and the merged variant and remove the others. To obtain the optimal overlap threshold, similar to [24], we test various thresholds for the *Qi overlap* measure (**Equation 7**) and choose the threshold which gives the highest F-similarity-based Maximal Matching F-score (FMMF).

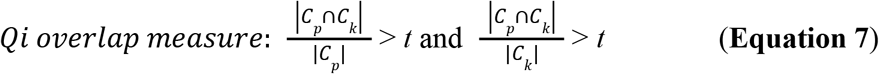

Here, *t* is the user-specified overlap threshold, *C*_*p*_ is a predicted complex and *C*_*k*_ is a known complex.

To gauge the accuracy of the RL algorithm, we use different evaluation measures to compare learned complexes with known complexes. The learned complexes are compared with the known complexes after removing nodes missing in the set of known complexes. We employ a variety of evaluation measures such as the FMMF, Community-wise Maximum F-similarity-based F-score (CMMF), and Unbiased Sn-PPV Accuracy (UnSPA) defined in Super.Complex [24], in addition to the Qi et al F-score [22], F-grand k-clique and F-weighted k-clique [6].

As discussed in [24], one of the more accurate methods for comparison is the FMMF which combines an adapted Maximal Matching Ratio (MMR) with its corresponding precision. The adapted MMR is computed as the fraction of known complexes matching predicted complexes, where the matches are computed as the sum of edge weights of a maximal matching in a bipartite graph between learned and known complexes. The edge weights used in the graph, matching learned and known complexes, are the F-similarity scores which are computed as follows.

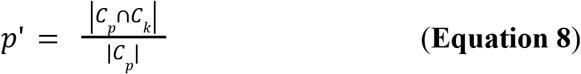

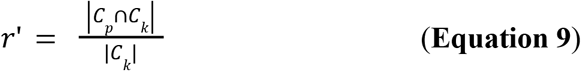

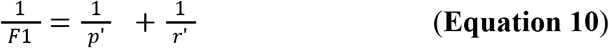

Here, *C*_*p*_ is a predicted complex and *C*_*k*_ is a known complex.

#### Experimental details - synthetic and protein interaction datasets

We first test the RL algorithm on a synthetic toy network **(Figure 3)** with 62 nodes and 78 edges, comprising 14 complexes. Of these 14 complexes, 7 are used as training complexes and 7 are used as testing complexes. The testing and training evaluation results can be found in **Table S1**N in the supplement.

**Figure 3.**
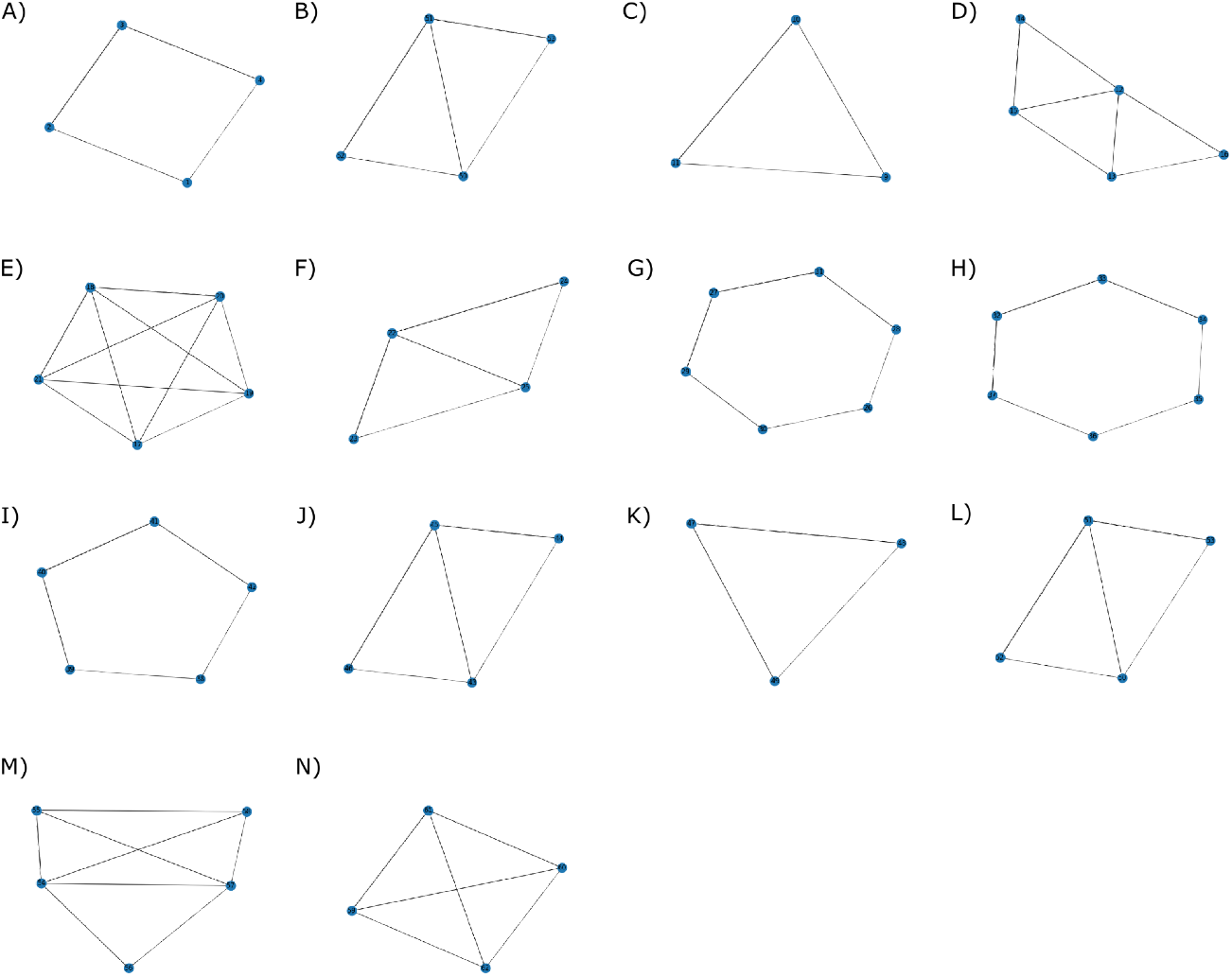
A synthetic disconnected toy network of complexes. Complexes A-G are used for training and H-N are used for testing the RL algorithm.

Next, we apply the RL algorithm on a human protein interaction network (hu.MAP 1.0 [6]) comprising 7778 nodes and 56,712 edges to learn candidate complexes using 188 known complexes; these complexes are obtained by pre-processing the CORUM protein complex database [26]. The pre-processing primarily involves discarding complexes that are internally disconnected or have fewer than 3 nodes, and merging complexes with a pairwise overlap of more than 0.6 Jaccard coefficient (see methods section of Super.Complex [24]). For a perfect comparison, we use the same pre-processing steps as in Super.Complex for both the network and complexes, as well as the same training and testing complexes (all input data was obtained from Super.Complex input data [24]). Since we only use the density feature in our algorithm, we also run the Super.Complex pipeline with only the density feature. The testing and training evaluation results can be found in **Table S2** in the supplement.

The RL algorithm is also tested on hu.MAP 2.0 [7], a human PPI network consisting of 10,433 nodes and 43,581 edges, obtained by considering only the edges with a weight of at least 0.02. Again, to compare with Super.Complex, we use the same training and testing complexes from hu.MAP 1.0, and the same preprocessing steps. We perform two experiments; in the first experiment, we transfer the value function trained on hu.MAP 1.0 and in the second, we train a new value function on hu.MAP 2.0. The testing and training evaluation results for hu.MAP 2.0 can be found in **Table S3** in the supplement.

## Results and discussion

### The value function converges in the training phase

During the training phase, we track the value for each encountered state (density) over time. Once the value of each of the states starts to converge, it can be assumed that the value has reached its optimum. **Figure 4** demonstrates the successful convergence of the value for each density.

**Figure 4.**
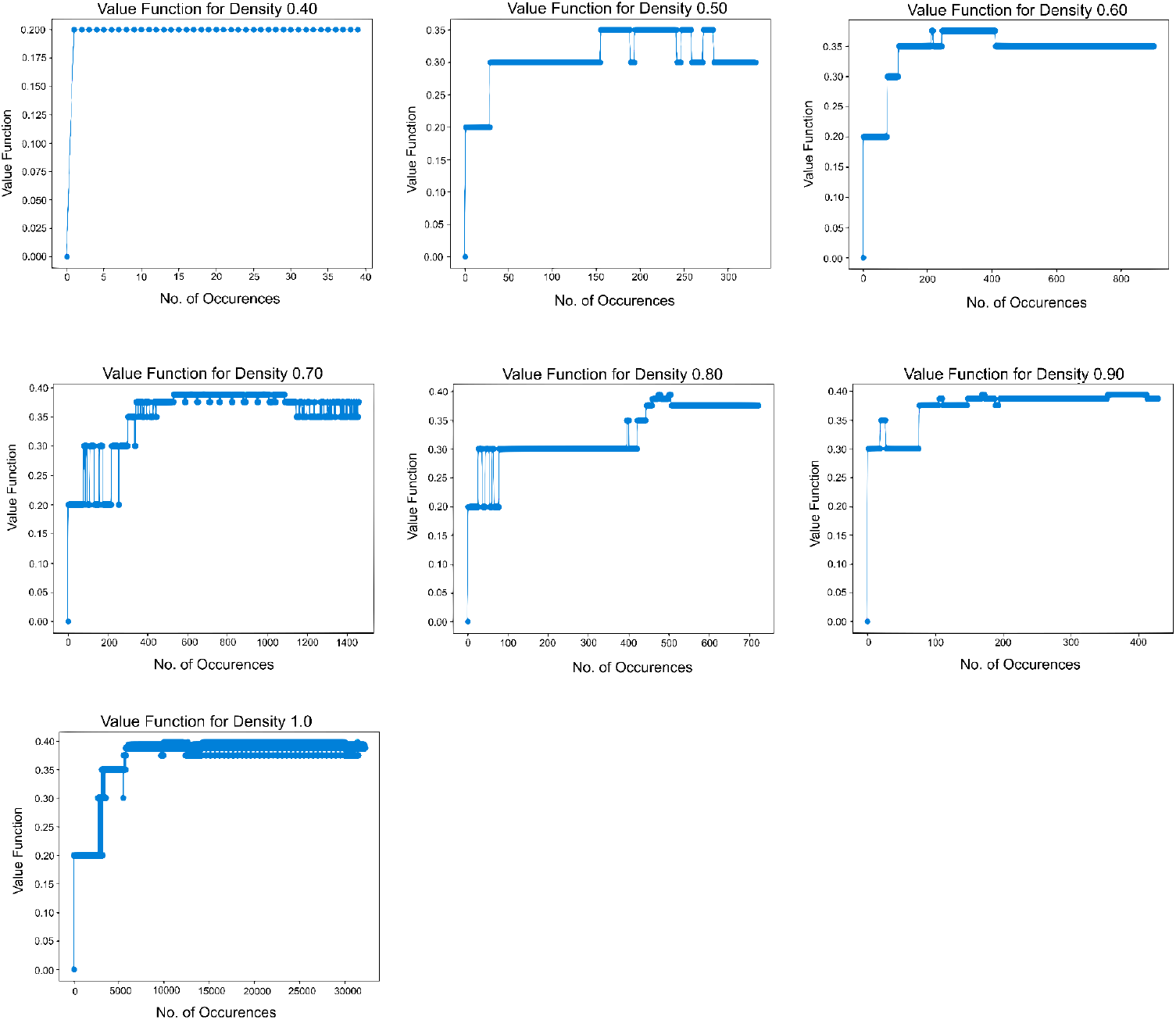
Convergence of the value for each density. Each density (0.4 through 1) encountered in the training for hu.MAP 1.0 is plotted to see how its value updates over iterations. Although the values fluctuate initially, they converge eventually indicating successful training.

We also investigate the relationship between a state’s density and its value. **Figure 5** shows that in the path to a final complex, subgraphs of higher densities are favored since they have higher values.

**Figure 5.**
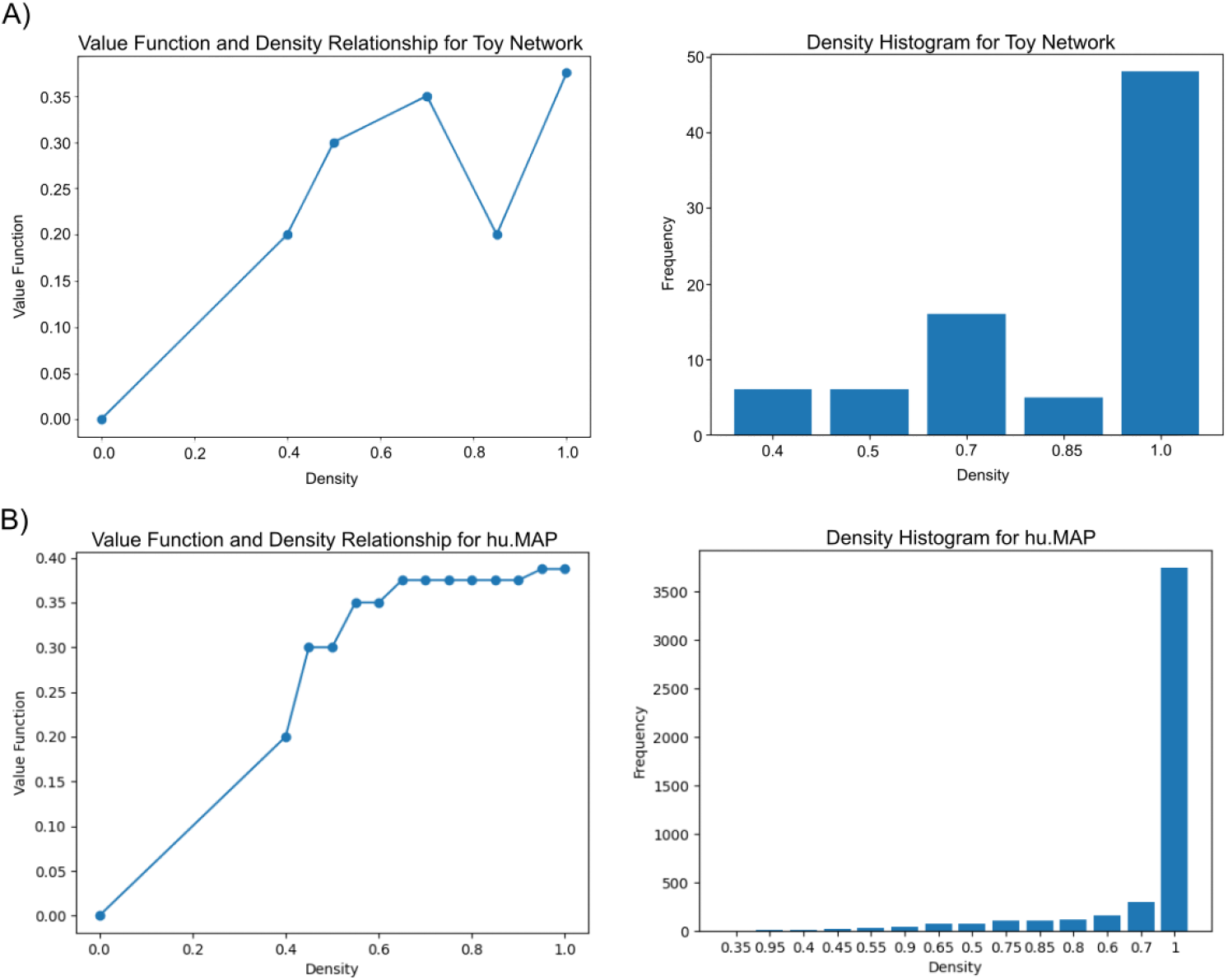
Higher values favor higher densities. **(A)** The graph on the toy network shows a positive correlation between value function and density and therefore, complexes and trajectories with higher densities are favored. The density histogram for the training phase shows a higher frequency for higher density subgraphs. **(B)** For hu.MAP 1.0, the graph shows a stronger correlation between density and value function. The density histogram again shows that higher densities are more frequently observed.

The learned value functions for the synthetic dataset and hu.MAP 1.0 enable us to accurately predict complexes on the respective networks, as shown in the next section. Further, employing transfer learning, we use the value function learned on hu.MAP 1.0 to accurately predict complexes on hu.MAP 2.0 (**Table S3**). This demonstrates that the value functions learned by the RL algorithm can be transferred for community detection problems on similar networks. We also directly train a value function on hu.MAP 2.0 using the same training complexes, and find that predicting complexes on hu.MAP 2.0 with this value function also gives accurate results (**Table S3**).

### The RL algorithm learns accurate communities on synthetic and real datasets

Recall the synthetic dataset containing 14 communities used to test the RL algorithm. The performance of the algorithm across different evaluation measures is excellent as summarized in **Table 1**.

**Table 1.** The RL algorithm has a strong performance on the synthetic toy dataset. The algorithm was trained on 7 toy complexes from a synthetic network of 62 nodes and 78 edges. It predicted 14 complexes which are evaluated against the 14 true complexes. Abbreviations: FMM, F-similarity-based Maximal Matching; CMMF, Community-wise Maximum F-similarity-based F-score; UnSPA, Unbiased Sn-PPV Accuracy; SPA, Sn-PPV Accuracy.

|  | FMM Precision | FMM Recall | FMM F-score | CMMF | UnSPA | Qi et al F1 score | SPA | F-Grand K-Clique | F-weighted K-Clique |
| --- | --- | --- | --- | --- | --- | --- | --- | --- | --- |
| **RL Algorithm** | 0.963 | 0.963 | 0.963 | 0.963 | 0.969 | 1.00 | 0.959 | 1.00 | 1.00 |

Next, we apply the algorithm on the real dataset, hu.MAP 1.0, by training it on 132 complexes. We test different Qi overlap thresholds (**Figure 6A**), in the RL algorithm, to merge highly overlapping complexes. The peak in **Figure 6A** occurring at 0.325 Qi overlap measure corresponds to the best FMMF score. For this value of the Qi overlap measure, the RL algorithm learns 1,157 complexes.

**Figure 6.**
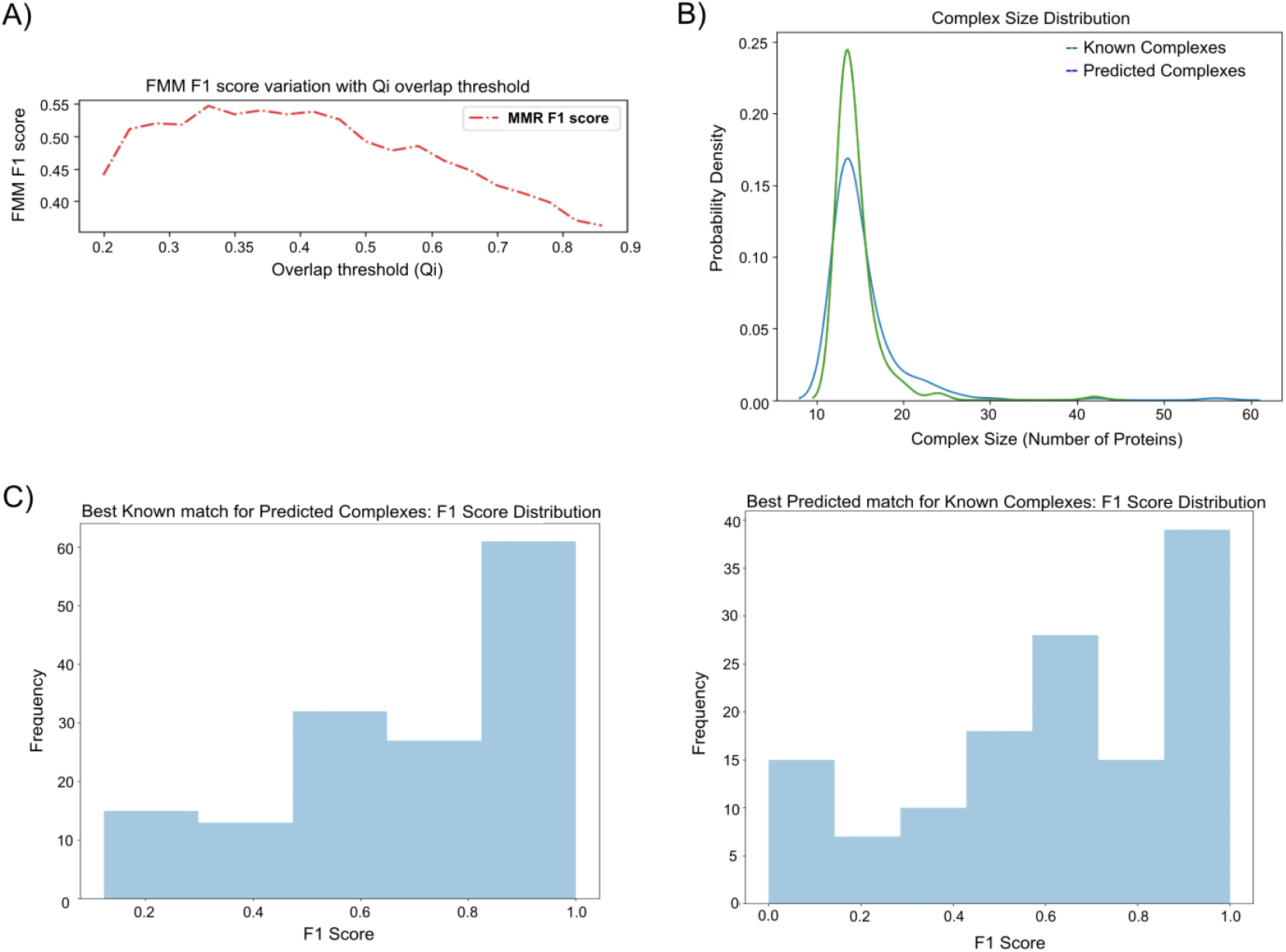
Evaluating the predictions of the RL algorithm on hu.MAP 1.0. **(A) The optimal Qi threshold is0.325**. We tested various overlap thresholds, *i.e*., Qi values (**Equation 7**) between 0.2 and 0.9 in 0.25 intervals. **Size distributions of known and predicted complexes**. This graph shows that the distribution of the sizes (no. of proteins) of the predicted and known complexes is very similar. (**C) F1 score distributions of the best-predicted match for known complexes and vice-versa**. In both cases, higher F1 scores have higher counts indicating accurate predictions.

For a perfect comparison, Super.Complex is tested on hu.MAP 1.0 using only the subgraph feature density. The best results from Super.Complex are obtained using a k-nearest neighbors classifier (with k = 76) to train a community fitness function, and from a search process for candidate complexes using maximal cliques as starting seeds and a pseudo-metropolis heuristic (with a probability of 0.1) for complex growth (with exploration probability ε 0.01). The candidate complexes are then merged with an overlap threshold of 0.2 Jaccard coefficient to yield 798 final complexes. In contrast, the RL algorithm predicts a higher number (1157) of complexes possibly resulting in a slightly higher (FMM) recall measure (**Table 2)**. The RL method achieves comparable performance to Super.Complex (**Table 2**), demonstrating the potential of applying reinforcement learning to community detection.

**Table 2.** The RL algorithm yields competitive accuracy compared to Super.Complex on hu.MAP 1.0. The learned complexes on hu.MAP 1.0 are evaluated against all the known cleaned CORUM complexes. Abbreviations: FMM, F-similarity-based Maximal Matching; CMMF, Community-wise Maximum F-similarity-based F-score; UnSPA, Unbiased Sn-PPV Accuracy; SPA, Sn-PPV Accuracy.

|  | FMM Precision | FMM Recall | FMM F-score | CMMF | UnSPA | Qi et al F1 score | SPA | F-Grand K-Clique | F-weighted K-Clique |
| --- | --- | --- | --- | --- | --- | --- | --- | --- | --- |
| **RL Algorithm** | 0.612 | 0.482 | 0.547 | 0.654 | 0.772 | 0.559 | 0.652 | 0.789 | 0.988 |
| **Super.Complex** | 0.835 | 0.457 | 0.591 | 0.720 | 0.803 | 0.658 | 0.692 | 0.991 | 0.999 |

Comparing **Tables 1** and **2**, we observe better accuracies for the algorithm on the toy dataset than those on hu.MAP 1.0. This could be attributed to the algorithm being better suited to finding non-overlapping communities, such as the toy communities, when compared to finding overlapping communities, such as the CORUM complexes on hu.MAP 1.0. Due to the current framework, the RL algorithm may not train well on overlapping communities because it trains on only one community in each episode, by giving a negative reward on adding a neighbor that does not belong to the community, even if it belongs to another overlapping community.

While accuracies are comparable, we note that the RL algorithm achieves faster running time relative to Super.Complex (**Table 3**). Specifically, the RL algorithm trains for ∼9s on one core of a personal computer (M1 chip @ 3.2 GHz), making the training significantly faster than Super.Complex’s training with the AutoML pipeline, which runs for ∼540s on 20 cores of a supercomputer (Intel(R) Xeon(R) CPU E5-2699 v3 @ 2.30GHz). Growing the candidate communities took ∼300s when running in parallel across 8 cores (3.2GHz) for the RL algorithm, compared to ∼20s when running in parallel across 72 cores (2.3GHz) for Super.Complex. This indicates that growing new complexes is also fast in the RL algorithm due to the simple inference using the value function lookup. We employ the 4 heuristics available in Super.Complex, with default parameters, to find the best one and perform a parameter sweep of 7 merge overlap thresholds for each heuristic. In total, we evaluate 28 heuristic-parameter combinations in Super.Complex; the same number of overlap thresholds used in the RL method. Overall, with the best parameters, the RL algorithm took ∼350s on the personal computer with 8 cores (note that only the prediction step is parallelized here), compared to Super.Complex which took ∼ 650s on a supercomputer with 72 cores (note that both the learning and the prediction step are parallelized here).

**Table 3.** The RL algorithm achieves a faster running time when compared to Super.Complex. The time reported here corresponds to runs using the best overlap threshold found for both methods and also using the best heuristic found in the case of Super.Complex.

| Method | Processor specifications | Training |  | Prediction |  | Post-processing |  | Total time (s) |
| --- | --- | --- | --- | --- | --- | --- | --- | --- |
|  |  | No. of cores | Time (s) | No. of cores | Time (s) | No. of cores | Time (s) |  |
| **RL Algorithm** | M1 chip<br>@ 3.2 GHz | 1 | 9 | 8 | 320 | 1 | 11 | 340 |
| **Super.Complex** | Intel(R) Xeon(R) E5-2699 v3<br>@ 2.30GHz | 20 | 540 | 72 | 17 | 1 | 112 | 669 |

The average time complexity of the prediction phase of the RL algorithm is*O*(*G*^*2*^*K*^*2*^*S/P*)where *G* is the average number of nodes in a complex, *K* is the average degree of the network and *S* is the number of seeds chosen (in our experiments, S is the number of nodes in the network). For Super.Complex with all subgraph features, the time complexity of the prediction phase is *O*(*XG*^*4*^*KS/P*). We note that the prediction phase of the RL algorithm scales better than that of Super.Complex. This is because the complexity of the subgraph feature extraction step reduces from *O(G^3^)*in Super.Complex to *O*(*GK*)in the RL algorithm; this reduction happens since the RL algorithm uses only the feature density with a constant model inference time (*X*). The time complexity of the RL training algorithm is *O*(*GK*)*T*, where *T* is the number of training complexes. In contrast, the training complexity of Super.Complex is *O*(*G*^*3*^*Tgpm/c*), where g is the number of generations, p is the population size, m is the number of machine learning models and feature preprocessor types tried, and c is the number of processes on the single compute node running the AutoML step.

We note that the RL algorithm can be very useful in community detection problems with a small number of known complexes, as demonstrated in our experiments (7 and 132 training complexes in the synthetic and real datasets). Even if the number of known complexes is small, for each complex, the value iteration procedure in the RL algorithm explores several trajectories to learn the complex, incidentally increasing the size of the training dataset used to learn the complexes. On the other hand, existing supervised community detection methods train on a dataset with a size equal to the number of training complexes. Other benefits of the RL algorithm include the lack of need for extensive hyperparameter tuning and the ability to predict complexes that do not contain smaller complexes. For comparison, in Super.Complex, at each stage of growth in a candidate complex, the pipeline seeks to yield a final protein complex, attempting 4 different heuristics, each with 1-2 hyperparameters. Contrastingly, the RL pipeline learns and traverses the optimal trajectory to find a complex without optimizing for intermediate complexes, and without the need for heuristics, thus saving on searching for parameters in the candidate complex growth step. Thus, the RL algorithm finds the best sequence of steps to grow a complex, while also being efficient.

In summary, relative to more sophisticated supervised ML strategies, the simplicity of the value iteration algorithm and the comparable accuracy along with improved efficiency demonstrates the great potential of the RL algorithm for solving community detection problems.

### The resulting clusters suggest functions for uncharacterized proteins

Importantly, the RL algorithm returns many well-known human protein complexes accurately (as would be expected from the precision measurements on withheld test complexes), several of which are illustrated in **Figure 7**. Notably, the algorithm also identifies several additional interaction partners and even potential new subunits within these systems, such as, for example, clustering the guanine nucleotide exchange factor RCC1L with proteins of the mitochondrial ribosome large subunit, consistent with a known role for RCC1L in mitochondrial ribosome biogenesis [27]. Similarly, the RL algorithm recapitulates the nutrient-response-related KICSTOR complex (SZT2, KPTN, ITFG2, and C12orf66) [28] on both hu.MAP 1.0 (SZT2, KPTN, ITFG2, TMCO4 and C12orf66) and hu.MAP 2.0 (SZT2, KPTN, ITFG2, TMCO4, BMT2, and KICS2), suggesting the uncharacterized transmembrane protein TMCO4 to be a potential new interaction partner, and it reconstructs the WAVE1/WAVE2 protein complexes, known regulators of actin filament and lamellipodia formation [29][30] while also suggesting involvement of KIAA1522, consistent with a recent suggestion for its involvement by Cho and colleagues [31]. In order to investigate the identified complexes interactively, visualizations are available for the 1157 learned complexes on the supporting website (see the Code and data availability section).

**Figure 7.**
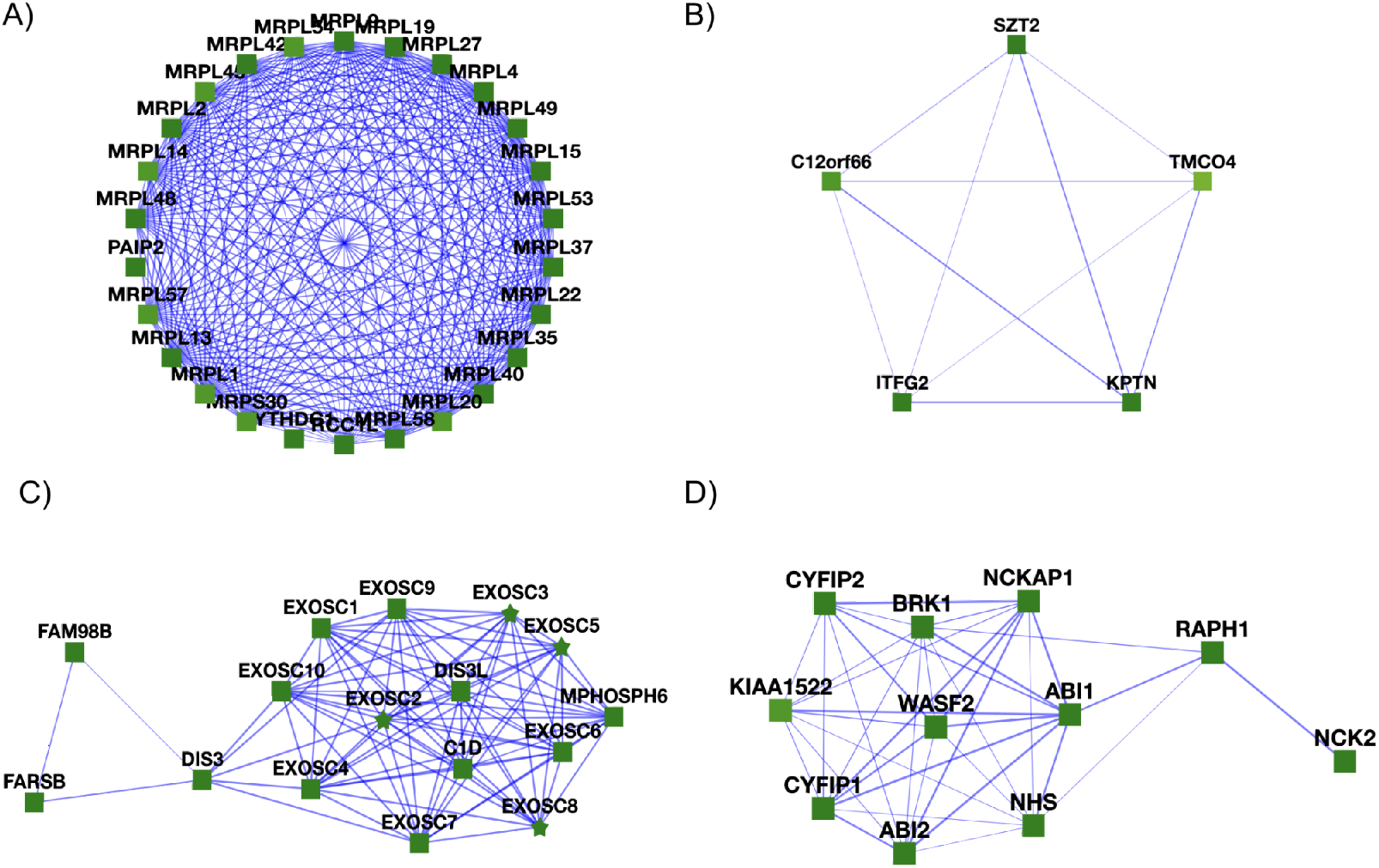
Learned complexes from the RL pipeline. **(A)** The large (39S) subunit of the mitochondrial ribosome is present in the RL-determined complexes, but broken up into multiple complexes, one of which is shown here. Note the presence of the guanine nucleotide exchange factor RCC1L, which is known to be essential for mitochondrial ribosome biogenesis [27]. **(B)** RL recapitulates the KICSTOR complex (C12orf66, KPTN, ITFG2, and SZT2), a multiprotein complex known to regulate mTORC1 and cells’ responses to available nutrient levels [28], but finds one additional putative subunit, the uncharacterized transmembrane protein TMCO4. **(C)** The exosome RNA processing complex is well-reconstructed by RL, with additional interactions observed to the tRNA synthetase FARSB and to FAM98B, a component of tRNA splicing ligase, consistent with possible associations among these systems [32]. **(D)** The WAVE1/WAVE2 protein complexes, known to regulate actin filament and lamellipodia formation [29][30], are reconstructed by RL, along with evidence for interaction with the uncharacterized protein KIAA1522. Notably, KIAA1522 was recently suggested by Cho and colleagues to bind WAVE and participate in a community of associated actin-organizing proteins [31].

Of particular interest are complexes corresponding to proteins with low annotation scores, as finding the proteins in complexes with better annotated proteins may help suggest potential functions for these otherwise minimally characterized proteins [33]. We searched specifically for such cases and highlighted complexes with uncharacterized proteins based on available UniProt annotations [34]. Some examples of learned complexes with uncharacterized or minimally characterized proteins are provided in **Figure 8**.

**Figure 8.**
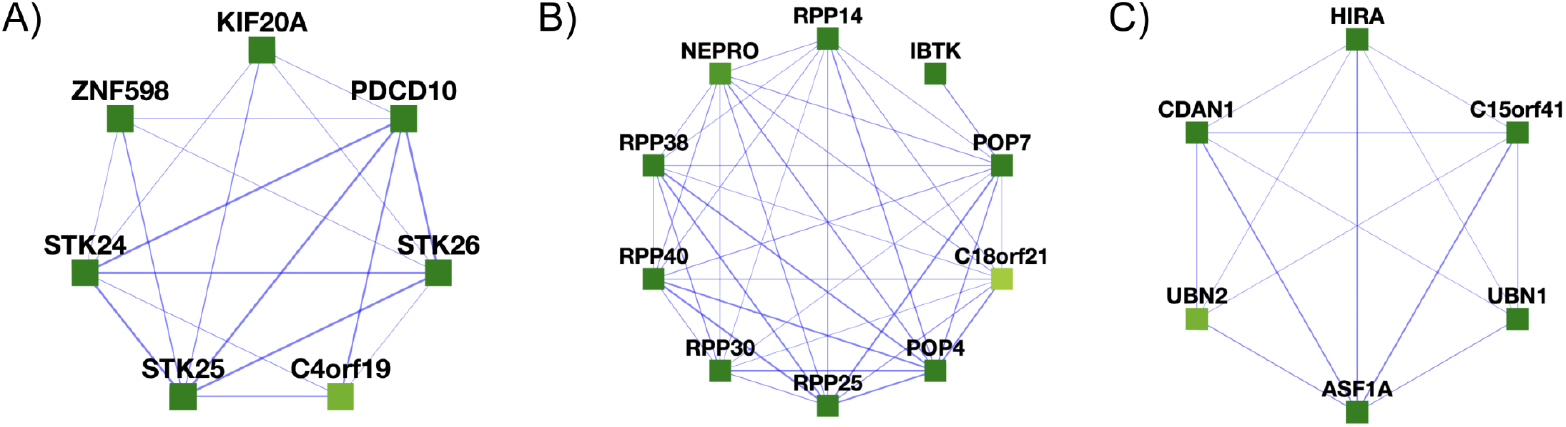
Participation in protein complexes by the uncharacterized proteins C4orf19 and C18orf21 and the minimally characterized protein C15orf41. **(A)** We find C4orf19 to belong to a larger complex composed of KIF20A, C4orf19, PDCD10, STK25, ZNF598, STK26, and STK24. **(B)** C18orf21 is found in a complex with 50% similarity to the Rnase/Mrp complex. **(C)** C15orf41 is found in a complex with 30% similarity to the cytosolic Codanin-1-Asf1-H3.1-histone H4–importin-4 complex.

For example, in **Figure 8A**, C4orf19 (chromosome 4 open reading frame 19) is broadly expressed across human cell types and tissues [35], with high protein levels in the kidney, liver, and GI tract [36][37] and while little is known about its function, an observed relationship between C4orf19 and colorectal cancer suggests that high expression levels might have some value as a marker for colorectal cancer [38], although elevated C4orf19 expression is also reported to show a favorable association with renal cancer survival [36][37]. Notably, four of the other proteins in this cluster (PDCD10, STK24, STK25, STK26) are known to associate into a complex with roles in maintaining epithelial integrity [39][40] and kidney water balance by regulating aquaporin trafficking and abundance in kidney tubule epithelial cells [41], suggesting a potential role for C4orf19 in normal kidney function. As for C4orf19, many of the proteins have been reported as potential biomarkers for bladder, gastric, pancreatic, and colorectal cancers [42]–[45].

As another example of a minimally characterized protein, C18orf21 (chromosome 18 open reading frame 21) is reported to possibly regulate the Rnase/Mrp complex, a ribonucleoprotein complex involved in RNA processing [46]. Both RL algorithm and Super.Complex concur on a connection for C18orf21 to RNA processing: from the learned complexes of Super.Complex, C18orf21 was found to be a part of a complex with a 50% overlap to the Rnase/Mrp complex, comprising all the proteins found in the RL algorithm’s learned complex (**Figure 8B**), adding support for this protein’s possible function in ribonuclease P RNA binding. Further, the RL algorithm learns a similar complex (C18orf21, IBTK, RPP30, POP4, and RPP25L) on hu.MAP 2.0 adding additional support to C18orf21’s function from the learned complexes on hu.MAP 1.0.

Somewhat more information can be gleaned for C15orf41 (chromosome 15 open reading frame 41), which, while minimally characterized, has recently been detected to interact with Codanin-1 (CDAN1) in human cells, and this interaction forms a tight, near stoichiometric complex [47]. Moreover, these studies reveal that mutation of C15orf41 can lead to the development of Congenital Dyserythropoietic type 1 disease (CDA-1) [47]. While its function is unknown, studies have noted a high sequence similarity between C15orf41 and archaeal Holliday junction resolvases, which are DNA repair enzymes that remove Holliday junctions [47], and it has been implicated in erythrocyte differentiation [48]. Its putative interaction partners within the complex (**Figure 8C**), HIRA and ASF1A, cooperate to promote chromatin assembly [49], and HIRA, ASF1A, and UBN1/2 form a complex and function in histone deposition of variant H3.3 into chromatin, independent of DNA replication [50]. CDAN1 and C15orf41 mutations lead to similar erythroid phenotypes and they were both eliminated from the same animal taxa, suggesting that these 2 proteins may participate in a shared pathway [51].

To obtain more support for overall physical association of proteins in this cluster, we modeled the 3D structure of the C15orf41-CDAN1 interaction using AlphaFold-Multimer [52], as implemented in Google Colab [53]. The AlphaFold model indicated a high-confidence interface spanning two distinct domains of CDAN1, one contributed from a domain spanning amino acids 1017-1203 and one covering one face of the larger N-terminal domain (2-997) centered on amino acids 427-472 and 843-997; these, in turn, interact with opposing surfaces of C15orf41 (**Figure 9**). The predicted structure is consistent with the prior experimental observation that the C-terminal 227 residues of CDAN1 (residues 1000-1227) are critical for the interaction [54]. To investigate the possibility of additional direct interactions between the C15orf41-CDAN1 heterodimer and one or more of the remaining proteins in the cluster, we took advantage of an available X-ray crystal structure that delineated the ASF1A interaction with HIRA residues amino acids 446-466 (PDB entry 2I32) [55] in order to further evaluate a larger complex. Using AF2-multimer, we modeled C15orf41, CDAN1 residues 2-74 and 286-1203 (omitting the intrinsically disordered segments, as determined by [53]), ASF1A residues 1-155 (omitting the intrinsically disordered tail), and HIRA residues 421-479, a somewhat larger segment known to be critical for the interaction with ASF1A [55]. As illustrated in full in **Figure 9**, AlphaFold suggested a binding site for ASF1A distinct from the C15orf41 binding site that, importantly, did not occlude the experimentally determined HIRA binding site, which AlphaFold also recapitulated. Thus, 3D structural modeling confirmed that 4 of the proteins in this cluster can be accommodated within the same overall multiprotein complex.

**Figure 9.**
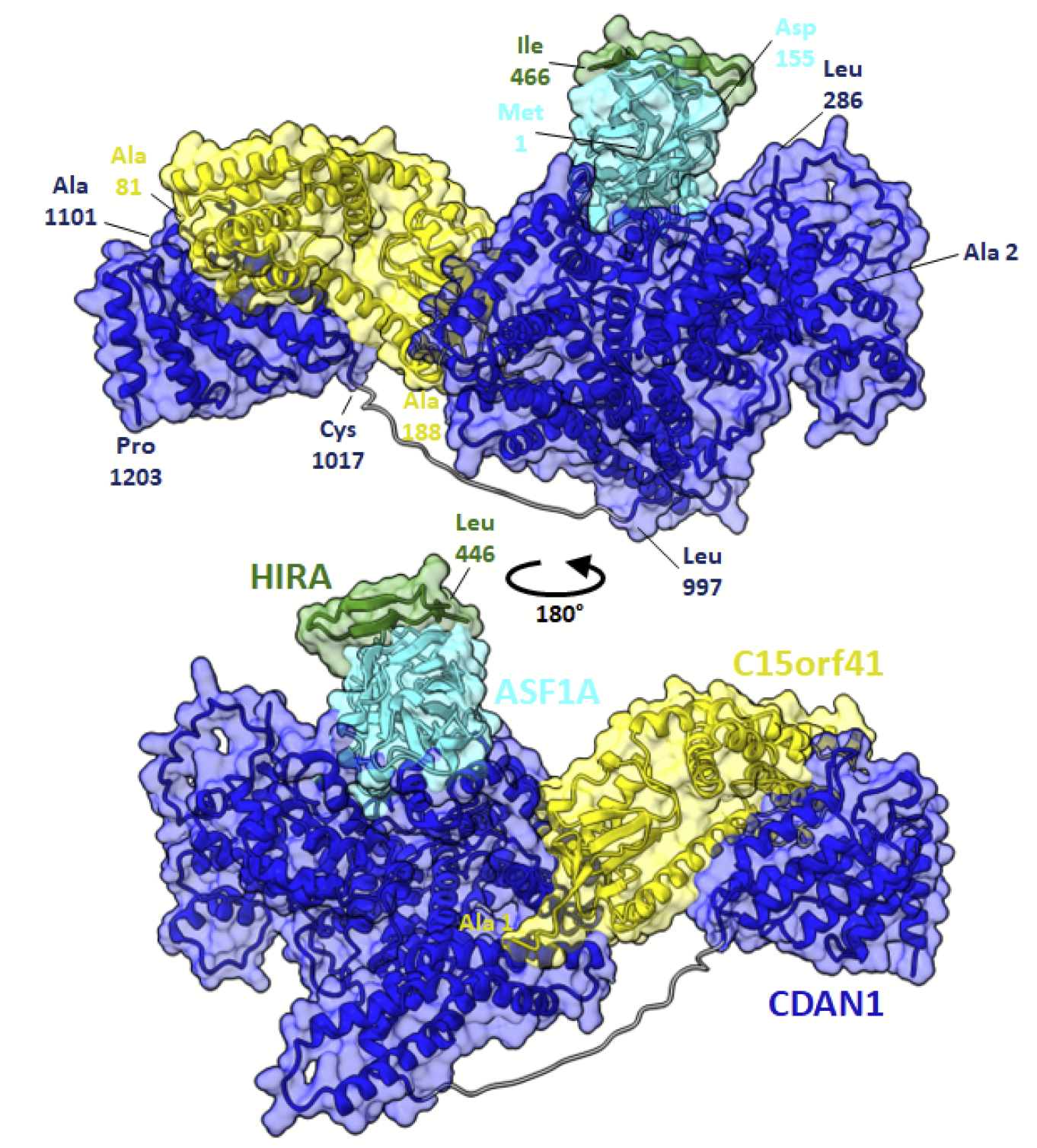
Structural modeling supports C15orf41, CDAN1, ASF1A, and HIRA participating in a large multiprotein complex. Using AlphaFold-multimer, we find that all 4 proteins can be simultaneously accommodated within a single multiprotein complex, here showing C15orf41’s modeled interaction with CDAN1 residues 2-74 and 286-1203, ASF1A residues 1-155, and HIRA residues 421-479. For illustration purposes, the known crystal structure of HIRA 446-466 [55] has been superimposed onto the AlphaFold model, which is available in full from the supporting GitHub repository with accompanying quality measurements.

Finally, C11orf42 was found as a subunit in a complex (C11orf42, SNX1, SNX5, VPS29, SNX2, COMMD9) that corresponds to a subcomplex of the Retromer or SNX/BAR complex (e.g. as in [56]), with supporting independent evidence from a learned complex from Super.Complex (C11orf42, SNX1, SNX5, and VPS29). This indicates that C11orf42 may be involved in trafficking with Retromer complex proteins, a notion supported by its localization to intracellular vesicles similar in nature to the other proteins in the complex [36][57]–[61]. Another example, C16orf91 constituted a complex (C16orf91, UQCC1, COX20, UQCC2) resembling a learned complex from Super.Complex (C16orf91, UQCC1, COX20).

## Conclusions and future work

In conclusion, we asked if reinforcement learning could be applied to learn to walk trajectories on a protein interaction network, and in this way more accurately determine protein complexes. Application of the method to currently available human protein interaction networks performed competitively with other algorithms, with comparable accuracy but a notable savings in computational time, and in turn led to clear predictions of protein function and interactions for several uncharacterized human proteins. We could support at least one of these, C15orf41, with independent evidence from 3D structural modeling.

Three main avenues can be explored to potentially improve the RL community detection algorithm, namely, improving the subgraph representations, the RL formulation, and the candidate community search process.

We currently represent a subgraph by a single feature, its density, which gives a problem formulation with a small state space. While performance may be negatively impacted, to improve accuracy, more subgraph features could be included in addition to density. Examples of other subgraph features that could be added include edge weight statistics, node clustering coefficient statistics, and degree correlation statistics. However, as the number of features increases, the state space increases exponentially. For instance, if we incorporate 18 topological features with a discretization of each feature into 10 bins, we increase the number of states to10. Different sample-based reinforcement learning methods could be applied to address this challenge and potentially give more accurate results.

The RL formulation could be modified to better accommodate overlapping community detection, by giving a positive reward if a neighbor is present in any of the training complexes, rather than only in the current complex being trained on. Note however that in this scenario, the reward given on choosing a neighbor changes dynamically if adding it in a future step no longer leads to a complex, due to the current subgraph building a different overlapping complex. Due to the dynamic rewards that need to be computed for each neighbor at each iteration, by checking whether the new subgraph is a part of any of the training complexes, computational time would increase significantly compared to the current static reward system, however, it may improve accuracy. Also, different rewards and discount factors can be experimented with in the training phase of the algorithm.

Finally, in the candidate community search process, there may be scenarios where there are multiple highest-scoring neighbors to add at an iteration in the growth process of a subgraph. Currently, we have only added one of the highest-scoring neighbors to grow the complex. Each of the other highest-scoring neighbors can be added as well to grow complexes we may have missed in the current method.

## Acknowledgments

This research was funded by grants to E.M.M. from the National Institute of General Medical Sciences (R35 GM122480), National Institute of Child Health and Human Development (R01 HD085901), and Welch Foundation (F-1515).

## Code and data availability

**Code, available on GitHub repository:**

https://github.com/marcottelab/RL_complex_detection

## Data

**All learned complexes from hu.MAP 1.0 with their corresponding scores:**

https://marcottelab.github.io/RL_humap_prediction/humap/res_pred_names.txt

**All learned complexes from hu.MAP 2.0 with their corresponding scores:**

https://marcottelab.github.io/RL_humap_prediction/humap2/res_pred_names_humap2.txt

**Interactive visualizations of results on hu.MAP 1.0 and hu.MAP 2.0:**

To investigate the complexes identified by the RL algorithm interactively, visualizations are available for the learned complexes on hu.MAP 1.0 and hu.MAP 2.0 on this website: https://marcottelab.github.io/RL_humap_prediction/

The website also provides the functionality of sorting complexes and proteins by their annotation score and the number of interactions with SARS-CoV-2 proteins [62].

**Structural model for the C15orf41, CDAN1, ASF1A, and HIRA heterotetramer, with sequences and assessments of model quality, can be downloaded from:** https://github.com/marcottelab/RL_humap_prediction/

(folder - CDIN1_CDAN1_2-74_286-1203_ASF1A_1-155_HIRA_421-479_1ModelRelaxed.zip)

## References

[1] A. L. Richards, M. Eckhardt, and N. J. Krogan, “Mass spectrometry-based protein–protein interaction networks for the study of human diseases,” Mol. Syst. Biol., vol. 17, no. 1, p. e8792, Jan. 2021, doi: 10.15252/msb.20188792.

[2] K. Titeca, I. Lemmens, J. Tavernier, and S. Eyckerman, “Discovering cellular protein-protein interactions: Technological strategies and opportunities,” Mass Spectrom. Rev., vol. 38, no. 1, pp. 79–111, Jan. 2019, doi: 10.1002/mas.21574.

[3] A. H. Smits and M. Vermeulen, “Characterizing Protein-Protein Interactions Using Mass Spectrometry: Challenges and Opportunities,” Trends Biotechnol., vol. 34, no. 10, pp. 825–834, Oct. 2016, doi: 10.1016/j.tibtech.2016.02.014.

[4] J. Snider, M. Kotlyar, P. Saraon, Z. Yao, I. Jurisica, and I. Stagljar, “Fundamentals of protein interaction network mapping,” Mol. Syst. Biol., vol. 11, no. 12, p. 848, Dec. 2015, doi: 10.15252/msb.20156351.

[5] T. M. Cafarelli, A. Desbuleux, Y. Wang, S. G. Choi, D. De Ridder, and M. Vidal, “Mapping, modeling, and characterization of protein-protein interactions on a proteomic scale,” Curr. Opin. Struct. Biol., vol. 44, pp. 201–210, Jun. 2017, doi: 10.1016/j.sbi.2017.05.003.

[6] K. Drew et al., “Integration of over 9,000 mass spectrometry experiments builds a global map of human protein complexes,” Mol. Syst. Biol., vol. 13, no. 6, p. 932, 08 2017, doi: 10.15252/msb.20167490.

[7] K. Drew, J. B. Wallingford, and E. M. Marcotte, “hu.MAP 2.0: integration of over 15,000 proteomic experiments builds a global compendium of human multiprotein assemblies,” Mol. Syst. Biol., vol. 17, no. 5, p. e10016, May 2021, doi: 10.15252/msb.202010016.

[8] A. Malovannaya et al., “Analysis of the Human Endogenous Coregulator Complexome,” Cell, vol. 145, no. 5, pp. 787–799, May 2011, doi: 10.1016/j.cell.2011.05.006.

[9] M. Y. Hein et al., “A Human Interactome in Three Quantitative Dimensions Organized by Stoichiometries and Abundances,” Cell, vol. 163, no. 3, pp. 712–723, Oct. 2015, doi: 10.1016/j.cell.2015.09.053.

[10] E. L. Huttlin et al., “The BioPlex Network: A Systematic Exploration of the Human Interactome,” Cell, vol. 162, no. 2, pp. 425–440, Jul. 2015, doi: 10.1016/j.cell.2015.06.043.

[11] E. L. Huttlin et al., “Architecture of the human interactome defines protein communities and disease networks,” Nature, vol. 545, no. 7655, Art. no. 7655, May 2017, doi: 10.1038/nature22366.

[12] C. Wan et al., “Panorama of ancient metazoan macromolecular complexes,” Nature, vol. 525, no. 7569, Art. no. 7569, Sep. 2015, doi: 10.1038/nature14877.

[13] K. J. Kirkwood, Y. Ahmad, M. Larance, and A. I. Lamond, “Characterization of Native Protein Complexes and Protein Isoform Variation Using Size-fractionation-based Quantitative Proteomics *,” Mol. Cell. Proteomics, vol. 12, no. 12, pp. 3851–3873, Dec. 2013, doi: 10.1074/mcp.M113.032367.

[14] A. R. Kristensen, J. Gsponer, and L. J. Foster, “A high-throughput approach for measuring temporal changes in the interactome,” Nat. Methods, vol. 9, no. 9, Art. no. 9, Sep. 2012, doi: 10.1038/nmeth.2131.

[15] P. C. Havugimana et al., “A Census of Human Soluble Protein Complexes,” Cell, vol. 150, no. 5, pp. 1068–1081, Aug. 2012, doi: 10.1016/j.cell.2012.08.011.

[16] M. A. Javed, M. S. Younis, S. Latif, J. Qadir, and A. Baig, “Community detection in networks: A multidisciplinary review,” J. Netw. Comput. Appl., vol. 108, pp. 87–111, Apr. 2018, doi: 10.1016/j.jnca.2018.02.011.

[17] G. D. Bader and C. W. Hogue, “An automated method for finding molecular complexes in large protein interaction networks,” BMC Bioinformatics, vol. 4, no. 1, p. 2, Jan. 2003, doi: 10.1186/1471-2105-4-2.

[18] G. Liu, L. Wong, and H. N. Chua, “Complex discovery from weighted PPI networks,” Bioinformatics, vol. 25, no. 15, pp. 1891–1897, Aug. 2009, doi: 10.1093/bioinformatics/btp311.

[19] M. Wu, X. Li, C.-K. Kwoh, and S.-K. Ng, “A core-attachment based method to detect protein complexes in PPI networks,” BMC Bioinformatics, vol. 10, no. 1, p. 169, Jun. 2009, doi: 10.1186/1471-2105-10-169.

[20] T. Nepusz, H. Yu, and A. Paccanaro, “Detecting overlapping protein complexes in protein-protein interaction networks,” Nat. Methods, vol. 9, no. 5, Art. no. 5, May 2012, doi: 10.1038/nmeth.1938.

[21] C. Lee, F. Reid, A. McDaid, and N. Hurley, “Detecting highly overlapping community structure by greedy clique expansion,” arXiv, 1002.1827, Jun. 2010. doi: 10.48550/arXiv.1002.1827.

[22] Y. Qi, F. Balem, C. Faloutsos, J. Klein-Seetharaman, and Z. Bar-Joseph, “Protein complex identification by supervised graph local clustering,” Bioinformatics, vol. 24, no. 13, pp. i250–i268, Jul. 2008, doi: 10.1093/bioinformatics/btn164.

[23] Y. Dong, Y. Sun, and C. Qin, “Predicting protein complexes using a supervised learning method combined with local structural information,” PLOS ONE, vol. 13, no. 3, p. e0194124, Mar. 2018, doi: 10.1371/journal.pone.0194124.

[24] M. V. Palukuri and E. M. Marcotte, “Super.Complex: A supervised machine learning pipeline for molecular complex detection in protein-interaction networks,” PLOS ONE, vol. 16, no. 12, p. e0262056, Dec. 2021, doi: 10.1371/journal.pone.0262056.

[25] E. C. Paim, A. L. C. Bazzan, and C. Chira, “Detecting Communities in Networks: a Decentralized Approach Based on Multiagent Reinforcement Learning,” in 2020 IEEE Symposium Series on Computational Intelligence (SSCI), Dec. 2020, pp. 2225–2232. doi: 10.1109/SSCI47803.2020.9308197.

[26] M. Giurgiu et al., “CORUM: the comprehensive resource of mammalian protein complexes-2019,” Nucleic Acids Res., vol. 47, no. D1, pp. D559–D563, Jan. 2019, doi: 10.1093/nar/gky973.

[27] J. D. Arroyo et al., “A Genome-wide CRISPR Death Screen Identifies Genes Essential for Oxidative Phosphorylation,” Cell Metab., vol. 24, no. 6, pp. 875–885, Dec. 2016, doi: 10.1016/j.cmet.2016.08.017.

[28] R. L. Wolfson et al., “KICSTOR recruits GATOR1 to the lysosome and is necessary for nutrients to regulate mTORC1,” Nature, vol. 543, no. 7645, Art. no. 7645, Mar. 2017, doi: 10.1038/nature21423.

[29] S. Suetsugu, H. Miki, and T. Takenawa, “Identification of two human WAVE/SCAR homologues as general actin regulatory molecules which associate with the Arp2/3 complex,” Biochem. Biophys. Res. Commun., vol. 260, no. 1, pp. 296–302, Jun. 1999, doi: 10.1006/bbrc.1999.0894.

[30] O. D. Weiner et al., “Hem-1 complexes are essential for Rac activation, actin polymerization, and myosin regulation during neutrophil chemotaxis,” PLoS Biol., vol. 4, no. 2, p. e38, Feb. 2006, doi: 10.1371/journal.pbio.0040038.

[31] N. H. Cho et al., “OpenCell: Endogenous tagging for the cartography of human cellular organization,” Science, vol. 375, no. 6585, p. eabi6983, Mar. 2022, doi: 10.1126/science.abi6983.

[32] D. Wichtowska, T. W. Turowski, and M. Boguta, “An interplay between transcription, processing, and degradation determines tRNA levels in yeast,” Wiley Interdiscip. Rev. RNA, vol. 4, no. 6, pp. 709–722, Dec. 2013, doi: 10.1002/wrna.1190.

[33] G. Kustatscher et al., “Understudied proteins: opportunities and challenges for functional proteomics,” Nat. Methods, pp. 1–6, May 2022, doi: 10.1038/s41592-022-01454-x.

[34] The UniProt Consortium, “UniProt: the universal protein knowledgebase in 2021,” Nucleic Acids Res., vol. 49, no. D1, pp. D480–D489, Jan. 2021, doi: 10.1093/nar/gkaa1100.

[35] “C4orf19 expression in human.” https://bgee.org/gene/ENSG00000154274 (accessed May 20, 2022).

[36] P. J. Thul et al., “A subcellular map of the human proteome,” Science, vol. 356, no. 6340, May 2017, doi: 10.1126/science.aal3321.

[37] “Tissue expression of C4orf19 - Summary - The Human Protein Atlas.” https://www.proteinatlas.org/ENSG00000154274-C4orf19/tissue (accessed Jun. 16, 2022).

[38] W. Wang et al., “Down-regulated C4orf19 confers poor prognosis in colon adenocarcinoma identified by gene co-expression network,” J. Cancer, vol. 13, no. 4, pp. 1145–1159, Jan. 2022, doi: 10.7150/jca.63635.

[39] X. Zheng et al., “CCM3 signaling through sterile 20-like kinases plays an essential role during zebrafish cardiovascular development and cerebral cavernous malformations,” J. Clin. Invest., vol. 120, no. 8, pp. 2795–2804, Aug. 2010, doi: 10.1172/JCI39679.

[40] M. Goudreault et al., “A PP2A phosphatase high density interaction network identifies a novel striatin-interacting phosphatase and kinase complex linked to the cerebral cavernous malformation 3 (CCM3) protein,” Mol. Cell. Proteomics MCP, vol. 8, no. 1, pp. 157–171, Jan. 2009, doi: 10.1074/mcp.M800266-MCP200.

[41] R. Wang et al., “Pdcd10-Stk24/25 complex controls kidney water reabsorption by regulating Aqp2 membrane targeting,” JCI Insight, vol. 6, no. 12, p. 142838, Jun. 2021, doi: 10.1172/jci.insight.142838.

[42] M. Xiong et al., “KIF20A promotes cellular malignant behavior and enhances resistance to chemotherapy in colorectal cancer through regulation of the JAK/STAT3 signaling pathway,” Aging, vol. 11, no. 24, pp. 11905–11921, Dec. 2019, doi: 10.18632/aging.102505.

[43] D. Stangel et al., “Kif20a inhibition reduces migration and invasion of pancreatic cancer cells,” J. Surg. Res., vol. 197, no. 1, pp. 91–100, Jul. 2015, doi: 10.1016/j.jss.2015.03.070.

[44] “PDCD10 programmed cell death 10 [Homo sapiens (human)] - Gene - NCBI.” https://www.ncbi.nlm.nih.gov/gene/11235 (accessed May 31, 2022).

[45] H.-P. Hsu, C.-Y. Wang, P.-Y. Hsieh, J.-H. Fang, and Y.-L. Chen, “Knockdown of serine/threonine-protein kinase 24 promotes tumorigenesis and myeloid-derived suppressor cell expansion in an orthotopic immunocompetent gastric cancer animal model,” J. Cancer, vol. 11, no. 1, pp. 213–228, Jan. 2020, doi: 10.7150/jca.35821.

[46] L. Liang, V. Chen, K. Zhu, X. Fan, X. Lu, and S. Lu, “Integrating data and knowledge to identify functional modules of genes: a multilayer approach,” BMC Bioinformatics, vol. 20, no. 1, p. 225, Dec. 2019, doi: 10.1186/s12859-019-2800-y.

[47] M. Shroff, A. Knebel, R. Toth, and J. Rouse, “A complex comprising C15ORF41 and Codanin-1: the products of two genes mutated in congenital dyserythropoietic anaemia type I (CDA-I),” Biochem. J., vol. 477, no. 10, pp. 1893–1905, May 2020, doi: 10.1042/BCJ20190944.

[48] R. Russo et al., “Characterization of Two Cases of Congenital Dyserythropoietic Anemia Type I Shed Light on the Uncharacterized C15orf41 Protein,” Front. Physiol., vol. 10, 2019, Accessed: Jun. 08, 2022. [Online]. Available: https://www.frontiersin.org/article/10.3389/fphys.2019.00621

[49] Y. Tang et al., “Structure of a human ASF1a/HIRA complex and insights into specificity of histone chaperone complex assembly,” Nat. Struct. Mol. Biol., vol. 13, no. 10, pp. 921–929, Oct. 2006, doi: 10.1038/nsmb1147.

[50] T. S. Rai et al., “Human CABIN1 Is a Functional Member of the Human HIRA/UBN1/ASF1a Histone H3.3 Chaperone Complex▿,” Mol. Cell. Biol., vol. 31, no. 19, pp. 4107–4118, Oct. 2011, doi: 10.1128/MCB.05546-11.

[51] G. Swickley et al., “Characterization of the interactions between Codanin-1 and C15Orf41, two proteins implicated in congenital dyserythropoietic anemia type I disease,” BMC Mol. Cell Biol., vol. 21, no. 1, p. 18, Mar. 2020, doi: 10.1186/s12860-020-00258-1.

[52] R. Evans et al., “Protein complex prediction with AlphaFold-Multimer.” bioRxiv, p. 2021.10.04.463034, Mar. 10, 2022. doi: 10.1101/2021.10.04.463034.

[53] M. Mirdita, K. Schütze, Y. Moriwaki, L. Heo, S. Ovchinnikov, and M. Steinegger, “ColabFold: making protein folding accessible to all,” Nat. Methods, vol. 19, no. 6, pp. 679–682, Jun. 2022, doi: 10.1038/s41592-022-01488-1.

[54] M. Shroff, A. Knebel, R. Toth, and J. Rouse, “A complex comprising C15ORF41 and Codanin-1: the products of two genes mutated in congenital dyserythropoietic anaemia type I (CDA-I),” Biochem. J., vol. 477, no. 10, pp. 1893–1905, May 2020, doi: 10.1042/BCJ20190944.

[55] Y. Tang et al., “Structure of a human ASF1a-HIRA complex and insights into specificity of histone chaperone complex assembly,” Nat. Struct. Mol. Biol., vol. 13, no. 10, pp. 921–929, Oct. 2006, doi: 10.1038/nsmb1147.

[56] T. Wassmer et al., “The retromer coat complex coordinates endosomal sorting and dynein-mediated transport, with carrier recognition by the trans-Golgi network,” Dev. Cell, vol. 17, no. 1, pp. 110–122, Jul. 2009, doi: 10.1016/j.devcel.2009.04.016.

[57] “Subcellular - C11orf42 - The Human Protein Atlas.” v21.http://proteinatlas.org/ENSG00000180878-C11orf42/subcellular.

[58] “Subcellular - SNX5 - The Human Protein Atlas.” v21.http://proteinatlas.org/ENSG00000089006-SNX5/subcellular.

[59] “Subcellular - VPS29 - The Human Protein Atlas.” v21.http://proteinatlas.org/ENSG00000111237-VPS29/subcellular.

[60] “Subcellular - SNX2 - The Human Protein Atlas.” v21.http://proteinatlas.org/ENSG00000205302-SNX2/subcellular.

[61] “Subcellular - SNX1 - The Human Protein Atlas.” v21.http://proteinatlas.org/ENSG00000028528-SNX1/subcellular.

[62] D. E. Gordon et al., “A SARS-CoV-2 protein interaction map reveals targets for drug repurposing,” Nature, vol. 583, no. 7816, Art. no. 7816, Jul. 2020, doi: 10.1038/s41586-020-2286-9.

