## Supplement for "Molecular complex detection in protein interaction networks through reinforcement learning"

### Supplementary figures

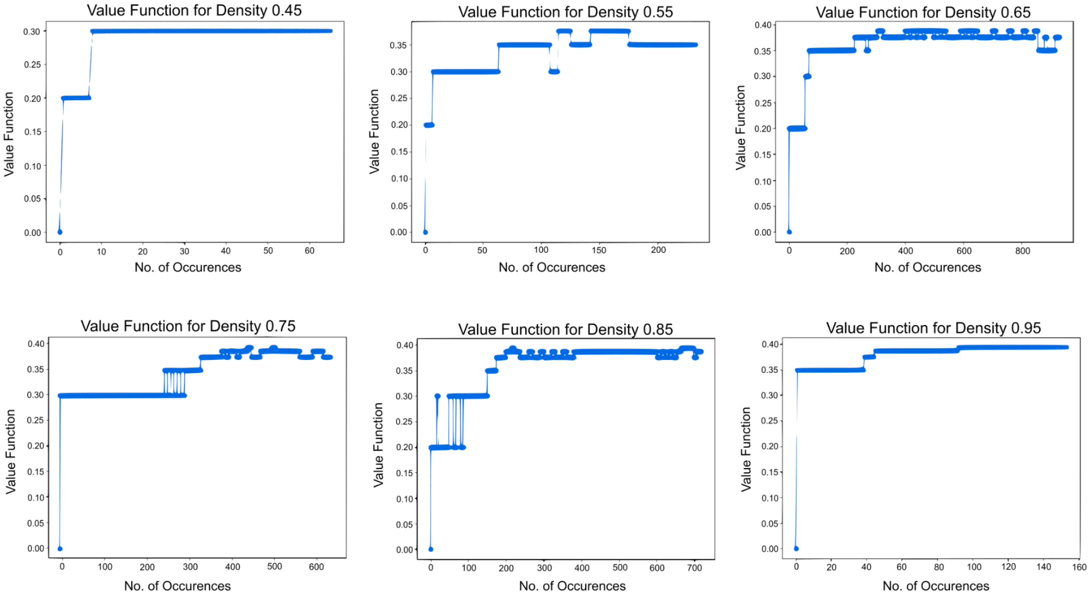


**Figure S1. Convergence of other scores from the training RL algorithm.** The values of the remaining densities encountered while training on hu.MAP 1.0 also converge eventually.

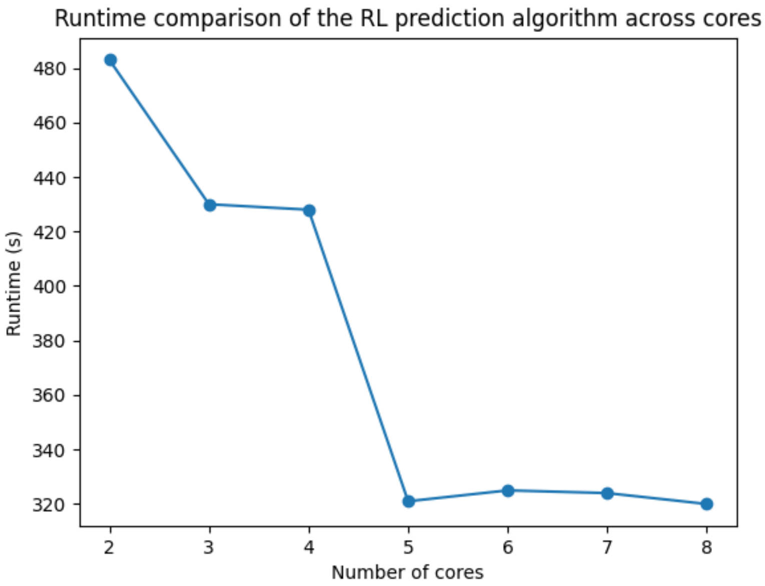


**Figure S2. As the number of cores used increases, the performance of the RL algorithm increases.** Here, the prediction algorithm was tested using 2 to 8 cores. The runtime decreases with an increasing number of processors ($P$), consistent with the average time complexity of the prediction algorithm scaling as $O(1/P)$.

### Supplementary tables

**Table S1. RL algorithm performance on training and testing toy complexes.** On the synthetic toy network, the RL algorithm predicted 14 toy complexes evaluated against each of the 7 training and 7 testing complexes.
Abbreviations: FMM, F-similarity-based Maximal Matching; CMMF, Community-wise Maximum F-similarity-based F-score; UnSPA, Unbiased Sn-PPV Accuracy; SPA, Sn-PPV Accuracy.

| | Evaluation set | FMM Precision | FMM Recall | FMM F-score | CMMF | UnSPA | Qi et al F1 score | SPA | F-Grand K-Clique | F-weighted K-Clique |
|---|---|---|---|---|---|---|---|---|---|---|
| **RL Algorithm** | Training | 0.971 | 0.971 | 0.971 | 0.971 | 0.976 | 1.00 | 0.952 | 1.00 | 1.00 |
| | Testing | 0.956 | 0.956 | 0.956 | 0.956 | 0.961 | 1.00 | 0.950 | 1.00 | 1.00 |

**Table S2. RL algorithm performance on training and testing hu.MAP 1.0 complexes.** The RL algorithm predicted 93 complexes with nodes from the training set, evaluated against the 132 training complexes, and 47 complexes with nodes from the testing set, evaluated against the 56 testing complexes. Super.Complex yields 56 and 49 complexes compared with the training and testing sets respectively after removing proteins absent in the known complexes.

| | Evaluation set | FMM Precision | FMM Recall | FMM F-score | CMMF | UnSPA | Qi et al F1 score | SPA | F-Grand K-Clique | F-weighted K-Clique |
|---|---|---|---|---|---|---|---|---|---|---|
| **RL Algorithm** | Training | 0.639 | 0.520 | 0.606 | 0.718 | 0.788 | 0.564 | 0.582 | 1.00 | 1.00 |
| | Testing | 0.642 | 0.538 | 0.618 | 0.586 | 0.702 | 0.552 | 0.500 | 0.993 | 0.996 |
| **Super.Complex (density)** | Training | 0.877 | 0.372 | 0.523 | 0.695 | 0.776 | 0.6 | 0.646 | 1.00 | 1.00 |
| | Testing | 0.821 | 0.704 | 0.758 | 0.789 | 0.856 | 0.779 | 0.811 | 0.985 | 0.999 |

**Table S3. RL algorithm performance on hu.MAP 2.0 complexes.**
**Row 1.** The RL algorithm uses the value function trained on hu.MAP 1.0 to predict complexes on hu.MAP 2.0 with an edge weight threshold of 0.1. This yields 1348 complexes, of which 85, 42, and 132 are compared with the training, testing, and all sets respectively after removing proteins absent in the known complexes. Parameters used include a Qi overlap threshold of 0.35.
**Row 2.** Super.Complex (using only feature density) uses the community fitness function trained on hu.MAP 1.0, to predict complexes on hu.MAP 2.0 with an edge weight threshold of 0.1. This yields 807 complexes, of which 56, 48, and 103 are compared with the training, testing, and both sets respectively after removing proteins absent in the known complexes. Parameters used include the pseudo-metropolis search heuristic starting with maximal cliques, merging overlaps with a Jaccard overlap threshold of 0.3.
**Row 3.** The RL algorithm (hu.MAP 2.0 trained) yields 2319 complexes, of which 81, 33, and 130 are compared with the training, testing, and both sets respectively after removing proteins absent in the known complexes. Parameters used include a Qi overlap threshold of 0.30.
**Row 4.** The union of the learned complexes by the RL algorithm on hu.MAP 2.0 with an edge weight threshold of 0.1 and 0.02 learned using the value function trained on hu.MAP 1.0. These are also the hu.MAP 2.0 complexes in the Data availability section. This yields 3614 complexes, of which 150, 53, and 247 are compared with the training, testing, and both sets respectively after removing proteins absent in known complexes. Parameters used include a Qi overlap threshold of 0.325.
**Row 5.** Super.Complex (all features), trained on hu.MAP 2.0 yields 582 complexes (after merging results from different edge weight thresholds), of which 55, 36 and 90 complexes are compared with the training, testing and both sets respectively after removing proteins absent in the known complexes. Parameters used include merging overlaps with a Jaccard overlap threshold of 0.1.

**Row 6.** 2 stage clustering from hu.MAP 2.0 yields 6948 complexes (after taking the union of results from different edge weight thresholds), of which 240, 143 and 385 complexes are compared with the training, testing and both sets respectively after removing proteins absent in the known complexes.

| No. | Method | No. of predicted complexes | Evaluation set | FMM Precision | FMM Recall | FMM F-score | CMMF | UnSPA | Qi et al F1 score | SPA | F-Grand K-Clique | F-weighted K-Clique |
|---|---|---|---|---|---|---|---|---|---|---|---|---|
| 1 | **RL Algorithm (hu.MAP 1.0 trained)** | 1348 | Training | 0.523 | 0.337 | 0.410 | 0.522 | 0.643 | 0.326 | 0.496 | 1.00 | 1.00 |
| | | | Testing | 0.623 | 0.467 | 0.534 | 0.563 | 0.665 | 0.542 | 0.643 | 1.00 | 1.00 |
| | | | All | 0.55 | 0.386 | 0.453 | 0.537 | 0.646 | 0.396 | 0.522 | 0.961 | 0.998 |
| 2 | **Super. Complex (density and hu.MAP 1.0 trained)** | 807 | Training | 0.824 | 0.374 | 0.515 | 0.664 | 0.748 | 0.628 | 0.625 | 1.00 | 1.00 |
| | | | Testing | 0.559 | 0.648 | 0.6 | 0.667 | 0.82 | 0.626 | 0.819 | 0.984 | 0.998 |
| | | | All | 0.681 | 0.449 | 0.541 | 0.657 | 0.774 | 0.625 | 0.693 | 0.991 | 0.999 |
| 3 | **RL Algorithm (hu.MAP 2.0 trained)** | 2319 | Training | 0.523 | 0.321 | 0.398 | 0.503 | 0.623 | 0.362 | 0.500 | 1.00 | 1.00 |
| | | | Testing | 0.406 | 0.239 | 0.301 | 0.333 | 0.454 | 0.247 | 0.491 | 1.00 | 1.00 |
| | | | All | 0.475 | 0.329 | 0.388 | 0.481 | 0.599 | 0.339 | 0.506 | 0.988 | 0.994 |
| 4 | **RL Algorithm (Union of 0.1 and 0.02 cutoff , hu.MAP 1.0 trained)** | 3614 | Training | 0.438 | 0.468 | 0.452 | 0.585 | 0.540 | 0.460 | 0.537 | 1.00 | 1.00 |
| | | | Testing | 0.349 | 0.493 | 0.409 | 0.542 | 0.477 | 0.455 | 0.732 | 1.00 | 1.00 |
| | | | All | 0.399 | 0.502 | 0.445 | 0.578 | 0.708 | 0.462 | 0.595 | 0.994 | 0.997 |
| 5 | **Super. Complex (all features and hu.MAP 2.0 trained)** | 582 | Training | 0.922 | 0.453 | 0.608 | 0.775 | 0.852 | 0.705 | 0.71 | 0.997 | 1.00 |
| | | | Testing | 0.908 | 0.696 | 0.788 | 0.824 | 0.855 | 0.846 | 0.798 | 0.984 | 0.998 |
| | | | All | 0.913 | 0.517 | 0.66 | 0.787 | 0.853 | 0.751 | 0.731 | 0.86 | 0.99 |
| 6 | **2 stage clustering (ClusterONE + MCL)** | 6948 | Training | 0.379 | 0.690 | 0.489 | 0.822 | 0.937 | 0.784 | 0.784 | 0.997 | 0.999 |
| | | | Testing | 0.335 | 0.856 | 0.482 | 0.898 | 0.931 | 0.905 | 0.938 | 0.764 | 0.959 |
| | | | All | 0.359 | 0.736 | 0.483 | 0.843 | 0.938 | 0.807 | 0.822 | 0.776 | 0.962 |
